# Distinct mechanisms by which antioxidant transcription factors Nrf1 and Nrf2 as drug targets contribute to the anticancer efficacy of Cisplatin on hepatoma cells

**DOI:** 10.1101/2023.08.16.553539

**Authors:** Reziyamu Wufuer, Keli Liu, Jingfeng, Meng Wang, Shaofan Hu, Feilong Chen, Shanshan Lin, Yiguo Zhang

## Abstract

Cisplatin (*cis*-Dichlorodiamineplatinum[II], CDDP) is generally accepted as a platinum-based alkylating agent type of the DNA-damaging anticancer drug, which is widely administrated in clinical treatment of many solid tumors. The pharmacological effect of CDDP is mainly achieved by replacing the chloride ion (*Cl^−^*) in its structure with H_2_O to form active substances with the strong electrophilic properties and then react with any nucleophilic molecules, primarily leading to genomic DNA damage and subsequent cell death. In this process, those target genes driven by the consensus electrophilic and/or antioxidant response elements (EpREs/AREs) in their promoter regions are also activated or repressed by CDDP. Thereby, we here examined the expression profiling of such genes regulated by two principal antioxidant transcription factors Nrf1 and Nrf2 (both encoded by *Nfe2l1* and *Nfe2l2,* respectively) in diverse cellular signaling responses to this intervention. The results demonstrated distinct cellular metabolisms, molecular pathways and signaling response mechanisms by which Nrf1 and Nrf2 as the drug targets differentially contribute to the anticancer efficacy of CDDP on hepatoma cells and xenograft tumor mice. Interestingly, the role of Nrf1, rather than Nrf2, is required for the anticancer effect of CDDP, to suppress malignant behavior of HepG2 cells by differentially monitoring multi-hierarchical signaling to gene regulatory networks. To our surprise, it was found there exists a closer relationship of Nrf1α than Nrf2 with DNA repair, but the hyperactive Nrf2 in *Nrf1α^−/−^*cells manifests a strong correlation with its resistance to CDDP, albeit their mechanistic details remain elusive.

## 1. Introduction

The mouse *Nfe2l1* gene (encoding NFE2-like CNC-bZIP transcription factor Nrf1) is situated at the distal end of chromosome 11, while human *Nfe2l1* gene is located on chromosome 17q21.3. The human prototypical Nrf1 is composed of 742 amino acids (aa), with a typical basic region-leucine zipper domain enabling transcriptional regulation of its cognate target genes by its physical interaction with NF-E2-binding sites [1], homologous to the *cis*-regulatory consensus electrophilic or antioxidant response elements (EpREs/AREs) in the promoter regions [2]. Subsequently, its long subtype called TCF11 was discovered to comprise 772 aa, with almost identical amino acid sequences and structural domains shared with Nrf1, except for an intrinsic loss of the Neh4L domain in the latter Nrf1 (Figure 1A) [3, 4]. Recently, the functional relationship between Nrf1 and TCF11 was further elucidated by their knockout and overexpression experiments in distinct cell lines, including human hepatoma (HepG2) cells [5]. The results demonstrated that TCF11 has exerted a significant inhibitory effect on the malignant proliferation of liver cancer and its migrating ability in the nude mouse xenograft model, which is much stronger than the effect of Nrf1. Conversely, the tumor proliferation and malgrowth were further augmented by specific knockout of Nrf1 and TCF11, albeit the latter TCF11 is, *de facto,* very lowly expressed in liver cancer [6, 7]. Therefore, it is inferable that Nrf1 and/or TCF11 have a potential target efficacy in preventing the malignant progression of liver cancer.

**Figure 1.**
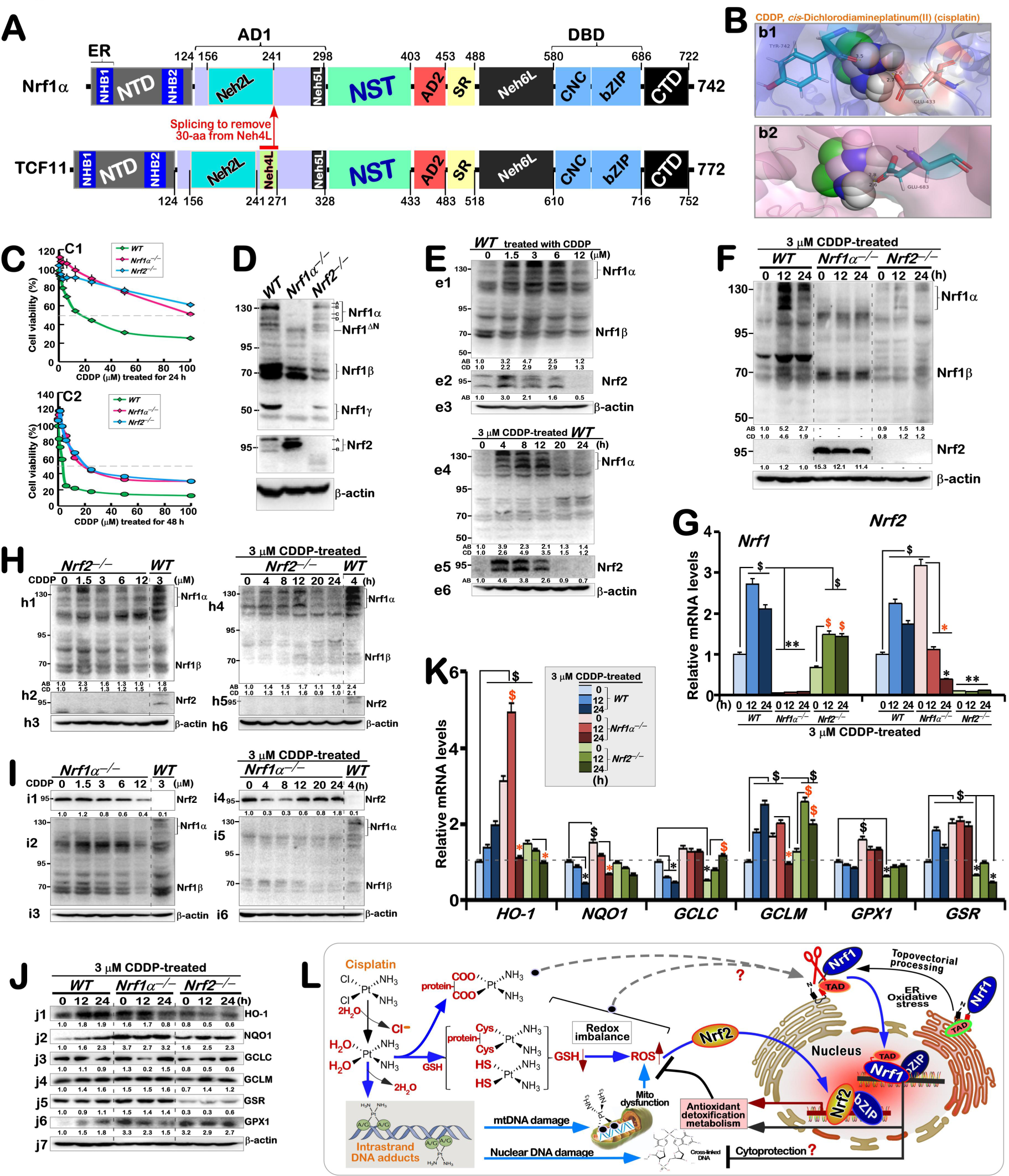
Distinct stimulation of Nrf1, Nrf2 and co-target genes by CDDP in different genotypic cell lines. **(A)** Structural schematics of Nrf1 and TCF11. Nrf1 has a loss of 30 amino acids within the Neh4L domain of TCF11, but the remaining structural domains of Nrf1 and its functional activities seem to be largely consistent with those of TCF11. **(B)** Two putative pockets of Nrf1/TCF11 are predicted to possess a strong binding activity with CDDP. ***(b1)*** When the coordinates of the active binding pocket were predicted to be 20.5, 3.5, −4.0 (x, y, z) (Detail in Figure S1. A) within Nrf1/TCF11 being docked with CDDP, with a binding energy of −7.64, this ligand is docked to E^433^ of TCF11 or E^403^ of Nrf1, with 3.5 or 2.7 of their O-H bond lengths, and Y^742^ (with 3.5 of the H-O bond length), respectively. ***(b2)*** When the predicted coordinates of the activity binding pocket are −11.5, 5.6, 15.5 (x, y, z) (Detail in Figure S1. A) within Nrf1/TCF11 being docked with CDDP, with another binding energy of −7.40, this ligand is docked to E^683^ of TCF11 or E^653^ of Nrf1, within different H-O bond lengths of 2.6 or 2.8, respectively. **(C)** Changes in the viability of distinct genotypic cell lines (*WT*, *Nrf1α^−/−^*, *Nrf2^−/−^*), that had been intervened for 24 h ***(c1)*** or 48 h ***(c2)*** with different doses of CDDP. The data were calculated as described elsewhere in the section of ‘Method and materials’. **(D)** Both Nrf1 and Nrf2 was identified by Western blotting of *WT, Nrf1α^−/−^* and *Nrf2^−/−^* cell lines. β-actin served as a loading control. **(E)** Western blotting analysis of the inducible expression levels of Nrf1 and Nrf2 in *WT* cells, that were treated with distinct doses of CDDP (*upper three panels*) for distinct lengths of time (*lower three panels*). **(F,G)** Distinct expression of Nrf1 and Nrf2 at protein and mRNA levels in *WT, Nrf1α^−/−^*, and *Nrf2^−/−^* cell lines that had been intervened with 3 μM CDDP for 0, 12 and 24 h, which were determined by Western blotting (***F***) and Real-time qPCR (***G***), respectively. **(H,I)** Western blotting analysis of Nrf1 and Nrf2 in both *Nrf2^−/−^*(***H***) and *Nrf1α^−/−^* (***I***) cell lines that had been treated with CDDP at different doses (*left panels*) for distinct lengths of time (*right panels*). In addition, some *WT* samples served as internal controls. **(J,K)** Differential expression levels of the indicated redox genes regulated by Nrf1 and/or Nrf2 were determined by Western blotting (***J***) and by Real-time qPCR (***K***), respectively, after *WT, Nrf1α^−/−^* and *Nrf2^−/−^* cell lines had been intervened or not with 3 μM CDDP. **(L)** A proposed model to give a better explanation of the putative mechanisms by which CDDP stimulates differential expression of Nrf1, Nrf2 and their target genes. The hydrolytic products of CDDP entering cells are prone to interaction with the thiol-based groups of proteins and/or the *N*-donating groups of nucleic acids to possibly damaging adducts and/or cross-linkages with target macromolecules. Consequently, such stress response, redox signaling, gene transcription, protein translation, and other relevant processes, particularly in which Nrf1 and Nrf2 are differentially implicated, are affected, and hence worthy to be further studied. In addition, it is notable that the intensity of all the above-described immunoblots was also quantified by the Quantity One 4.5.2 software as shown below each of indicated protein bands. Those real-time qPCR data were shown as fold changes (mean ± SD, n = 3 × 3), which are representative of at least three independent experiments being each performed in triplicates. Significant increases ($, p < 0.05; $$, p < 0.01) and significant decreases (\**p* < 0.05; \*\**p* < 0.01) were statistically analyzed, when compared with their corresponding control values (measured at 0 h, i.e., *t0*), respectively. Some symbols ‘$ or *’ also indicate significant differences in the CDDP intervention of *Nrf1α^−/−^*or *Nrf2^−/−^* cell lines, relative to their respective controls at *t0*.

By contrast with Nrf1, another homologous member Nrf2 of this CNC-bZIP family is commonly accepted as a master regulator of transcription factors involved in antioxidant, detoxification and cytoprotective responses to oxidative and electrophilic stress [8–10]. However, Nrf1 rather than Nrf2 is, as a matter of fact, indispensable for maintaining cell homeostasis and organ integrity in the changing internal and external environments, albeit Nrf2 is also required for this process and involved in cell growth, metabolism, autophagy, aging and other aspects [2]. Such a striking distinction was unraveled by the study on the phylogenetic tree of this highly conserved CNC-bZIP family, demonstrating that the emergence of membrane-bound Nrf1 and closer homologues occurs at an earlier ancestral stage of their evolutional process from marine bacteria to humans, as compared with the water-soluble Nrf2 [11]. In fact, it was indeed found that the gene-targeting loss of Nrf1’s function in all experimental mice and cell lines leads to severe endogenous oxidative stress and also results in spontaneous development of different pathological phenotypes, resembling human nonalcoholic steatohepatitis (NASH) and ultimate hepatoma [2, 12]. In addition, Nrf1 also plays a critical role in embryonic development [13] and osteoblast formation [14], as well in the maintenance of brain and heart integrity [15, 16]. Collectively, these results indicate that Nrf1 fulfills certain key important biological functions in the cytoprotective defense during life process, in addition to participating in the essential redox regulation. Thereby, it is quite plausible that the exploration of drugs (or leading-compounds) targeting for Nrf1 (and TCF11) may provide the strong evidence for its impact on the malignant development of tumors. This starting point enables Nrf1 to be paved as a precision therapeutic target for cancer treatment, along with another certain reference value of cancer intervention in the preventive strategy.

In the past three decades, the overwhelming majority of relevant researches in this CNC-bZIP field had been focused disproportionately on Nrf2 to be developed as a putative drug target for cancer treatment [9, 17]. Rather, it is unexpectedly unveiled that Nrf2 has a potent capability to exert its ‘double-edged sword’ role in suppressing or promoting cancer [18–20]. Relative to Nrf2, certainly much less attentions have been paid on Nrf1, albeit the latter CNC-bZIP factor possesses a stronger potential to fulfill its unique indispensable role in cancer prevention and treatment [12, 21]. In our previous studies, it was found that the compounds *tert*-butylhydroquinone (tHBQ) [22]), dithiothreitol (DTT) [23] and tunicamycin (TU) [24] stimulate a certain inducible effect on transactivation of Nrf1, but the related research is limited only to elucidating the differential expression of Nrf1 and Nrf2 in distinct cellular responses to oxidative, reductive or proteotoxic stresses, respectively. The results indicated that under different conditions (set for the endogenous absence of Nrf1 or Nrf2, along with intervention with theexogenous inducer), these two factors can make their unique and cooperative contributions to the basic composition of antioxidant and detoxification genes (i.e.*, GCLC, GCLM, HO1, NQO1, SOD, CAT*) and/or to stimulated expression of these and other cytoprotective genes in their regulatory network. Of note, Nrf1 has another potent capability to govern the functional activity of Nrf2 (and its targets) insomuch as to be strictly confined to a certain extent, albeit itself transcriptional expression is also regulated by Nrf2.

In this study, cisplatin (also called *cis-diaminodichloroplatinum*[II], hence abbreviated as CDDP) was employed as an external stimulus to explore the regulation of putative Nrf1-transactivating drugs in redox defense system of distinct genotypic cell lines. Under the premise of such Nrf1-targeting cancer treatment, we aim at discovering a new function of CDDP as an old-chemotherapeutic drug in translational medicine. CDDP is clearly known as a platinum-based anticancer drug with its square planar structure, but its biological activity is endowed only within the *cis-*, but not *trans-,* structures [25, 26]. This compound was firstly synthesized by *M. Peyrone* [27] until the late 19^th^ century, when its chemical structure was elucidated by Alfred Werner [26]. Notably, it was only in the 1960s that CDDP was scientifically studied, but in the begin with initial observations by Rosenberg showing that those electrolytic products of some platinum mesh electrodes could inhibit the cell division and proliferation of *E. coli*, which led to the exploration of the research value of platinum in tumor chemotherapy [27]. Later, CDDP was approved by the Food and Drug Administration (FDA) for cancer treatment in 1978 and is still considered as one of the most powerful anticancer drugs used until now [28]. Also, it should be of a crucial importance to note that CDDP was unraveled to pass through cell membranes without carrier transport after its dissolution in the body, and also possesses dual functional groups similar to those alkylating agents [29]. Such pharmacological effects of CDDP are mainly achieved by replacing the chloride ion (*Cl^−^*) in its structure with H_2_O to form active substances with strong electrophilic properties and then react with any nucleophilic molecules, particularly those of cellular components. Thereby, this drug is enabled to generate various adducts and cross-linkages with DNA alone or plus proteins, leading to significant DNA damages in the nuclear and mitochondrial genomes, cell cycle arrest and apoptosis [26]. Besides, CDDP also causes mitochondrial dysfunction by increasing intracellular levels of reactive oxygen species (ROS) and subsequent lipid peroxidation [29]. In this process, Nrf1 and Nrf2 along with target genes driven by a battery of EpREs/AREs within their promoter regions are stimulated by CDDP. By examining the expression profiling of such genes regulated by Nrf1 and/or Nrf2, we have here demonstrated distinct molecular pathways, cellular metabolisms, signaling response mechanisms by which Nrf1 and Nrf2 as two drug targets are differentially or coordinately contributable to the anticancer efficacy of CDDP on the hepatoma (HepG2) cells and xenograft tumor mice. Interestingly, the role of Nrf1, rather than Nrf2, is required for such an anticancer effect of CDDP, to suppress the malignant behaviors of HepG2 cells by differentially monitoring multi-hierarchical signaling responses to a number of key gene regulatory networks. To our surprise, it was found that there exists a closer relationship of Nrf1α than Nrf2 with the DNA damage repair, but the hyperactive Nrf2 in *Nrf1α^−/−^* cells manifests a strong correlation with its resistance to CDDP, albeit their detailed mechanisms need to be further explored.

## 2. Materials and methods

### 2.1 Chemicals and antibodies

Cisplatin (CDDP) is a platinum-containing metal complex with formula of *Cl_2_H_6_N_2_Pt* and MW of 300.05 g/mol (CAS No. 15663-27-1, Aladdin, Shanghai, China). In this study, 7.50 mg of CDDP powder was completely dissolved to obtain a stock solution of 100 mM CDDP stored at −20°C (Follow the instructions on the purchase website: https://www.aladdin-e.com/zh_cn/d109812.html). The antibody against Nrf1 was made in our own laboratory. These antibodies against Nrf2 (ab62352), GCLC (ab207777), GCLM (ab126704), HO-1 (ab52947), GPX1 (ab108427), XBP1 (AB109221), ATF4 (ab184909), ATF6 (ab227830), P4HB (ab137180), JUK (ab208035), p-JNK (ab124956), p-P38 (ab178876), PTEN (ab32199) and PI3K (ab302958) were obtained by Abcam (Cambridge, UK). Such antibodies against GSR (D220726), NQO1 (D26104), HK1 (D221854), ACCα (D155300), FASN (D272601), SCD1 (D162163) and CPT2 (D194980) were purchased from Sangon Biotech (Shanghai, China). Antibodies against BIP (bs-1219R), Chop (bs-20669R), p-IRE1 (bs-16698R), CDH1 (bs-10009R) and P65 (bs-0465R) were from Bioss (Beijing, China). Those antibodies against to p-eIF2α (#5199) was from CST (Boston, USA), p-PERK (sc-32577) from Santa Crus (CA, USA). The other antibodies against TXN1 (A5894), GLUT1 (A5514), GLUT4 (A11208), ATF4 (A5514), HSP60 (A5629), FGF21 (A5710), ERK (A5029), p-ERK (A5036), MEK (A5606), p-MEK (A5191) and P38 (A5017) were obtained from bimake (Texas, USA). Additional antibodies against G6PD (A1537), HK2 (A0994), LDHA (A7637), PDH (A1146) were from ABclonal (Wuhan, China). The β-actin (TA-09) was obtained from ZSGB-BIO (Beijing, China).

### 2.2 Cell lines and cell culture

The human hepatocellular carcinoma HepG2 cells (i.e. *Nrf1/2^+/+^* or *WT*) were obtained from the American Type Culture Collection (ATCC, Manassas, VA, USA). Two derived cell lines with knockout of *Nrf1α^−/−^* or *Nrf2^−/−^* were established in our own laboratory, and their relevant characterization had been described elsewhere by Qiu *et al* [6] and Ren *et al* [30]. The fidelity of HepG2 cells had been conformed to be true by its authentication profiling and STR (short term repeat) typing map (Shanghai Biowing Applied Biotechnology Co., Ltd, Shanghai, China). All these cell lines were incubated in a 37°C with 5% CO2, and allowed for growth in DMEM with 25 mmol/L high glucose (Gibco, USA), 10% fetal bovine serum (Gibco, USA) and 100 units/mL penicillin-streptomycin (MACKLIN, Shanghai, China).

### 2.3 Cell proliferation and toxicity assays

Three distinct genotypic cell lines (*WT, Nrf1α^−/−^, Nrf2^−/−^*) were seeded at a density of 6 × 10^3^ cells in each of 96 well plates. After the cells completely adhered, they were treated with different concentrations of CDDP (0.75, 1.5, 3, 6, 12, 25, 50 or 100 μM) for 24 h or 48 h. Additionally, MTT reagents (10 μl per well of 10 mg/ml stocked) was used to detect by the cell viability. Each of these experiments was repeated in five separated wells.

### 2.4 Detection of ROS, cell cycle and apoptosis by flow cytometry

To measure intracellular ROS levels, equal numbers (3 × 10^5^ cells in each well of 6-well plates) of experimental cells (*WT, Nrf1α^−/−^, Nrf2^−/−^*) were allowed for 24-h growth. After reaching 80% of their confluence, the cells were treated with 3 μM CDDP for distinct time periods (i.e., 0, 12, 24 h) before being collected. The intracellular ROS were determined by flow cytometry, according to the instruction of a ROS assay kit (S0033S, Beyotime, Shanghai, China). The obtained data are presented by column charts at a ratio of *Ex/Ex_WT0_*. Next, experimental cells were treated with 3 μM CDDP for 24 h and subjected to measuring cell cycle or apoptosis according to the instructions of relevant kits (C1052 or C1062S, Beyotime, Shanghai, China), respectively. The resulting data are presented by column charts of a percentage or a ratio of *Ex/Ex_WT0_*.

### 2.5 Detection of total antioxidant capacity (T-AOC), NADP^+^/NADPH *and* glucose

Each of experimental cell lines (*WT, Nrf1α^−/−^, Nrf2^−/−^*) was seeded in 5-cm diameter dish (6×10^5^ cells/dish) and allowed for growth before being treated with 3 μM CDDP for 0, 12 or 24 h and collected by centrifuging at 1000 × *g*. Subsequently, the total antioxidant capacity (T-AOC) of all cell samples was determined by the ABTS method, according to the instruction of the capacity detection kit (A015-2-1, Jiancheng Bioengineering Institute, Nanjing, China). The obtained data are presented in the form of a ratio column chart (*Ex/Ex_WT_0*). Next, CDDP-treated cells was incubated with 200μL of NADP^+^/NADPH extraction buffer and then collected by centrifuging at 12000 × *g* for 5-10 min at 4°C. The supernatant of cell lysates was saved and then subjected to measuring the NADP^+^/NADPH ratio, according to the relevant kit instructions (S0179, Beyotime, Shanghai, China). Lastly, CDDP-treated cells were washed twice with PBS and incubated with 200 μL of lysis buffer before being collected by centrifuging at 12000 × *g* for 5 min. The intracellular glucose content was tested according to the instructions of the reagent kit (S0201S, Beyotime, Shanghai, China).

### 2.6 Analysis of key gene expression by real-time quantitative PCR

All experiment cells (3 × 10^5^ per well) were seeded in 6-well plates and then allowed for growth to reach 80% of their confluence, before they were treated with 3 μM CDDP for 12 h or 24 h. The total RNAs were extracted and subsequently subjected to reaction with a reverse transcriptase to synthesize the first strand of cDNAs. This cDNAs were used as the template of real-time quantitative PCR (RT-qPCR) with each pair of the indicated gene primers (as listed in supplementary Table 1) to determine the mRNA expression levels of those genes in different genotypic cell lines (*WT, Nrf1α^−/−^, Nrf2^−/−^*). All these experiments were carried out within the Go Taq real-time PCR detection system by a CFX96 instrument. The resulting data were analyzed by the Bio-Rad CFX Manager 3.1 software and then presented graphically.

### 2.7 Analysis of key protein expression *by* western blotting

Experimental cell lines (*WT, Nrf1α^−/−^, Nrf2^−/−^*) were cultured at 6-well plates (4 × 10^5^ cells per well) and allowed for exposure into the indicated experimental settings. All those examined protein expression abundances were determined by Western blotting with their specific antibodies. Briefly, after quantitating total proteins in each of samples with the BCA protein reagent (P1513-1, ApplyGen, Beijing, China), they were separated by SDS-PAGE gels containing 8% to 12% polyacrylamide and then transferred onto the PVDF membranes. The membranes were blocked in Tris-buffered saline containing 5% non-fat dry milk at room temperature for 1 h, and incubated with each of specific primary antibodies overnight at 4°C. The antibody-blotted membranes were washed with PBST three times, before being re-incubated with the secondary antibodies at room temperatures for 2 h and then subjected to the imaging of immunoblots by Bio-Rad instrument. The intensity of immunoblotted proteins was calculated by the Quantity One software, while β-actin was used as a loading control.

### 2.8 Immunocytochemistry of γ-H2AX by confocal microscopy

Each group of cell lines (3 × 10^5^) were allowed for growth in each well of 6-well plates. After the cells were fully adhered, they were treated with 3 μM CDDP for different lengths of time (0 or 24 h) and then washed with PBS, before being tested for its DNA damage. The fixed cells were blocked and then incubated with the antibody against γ-H2AX (C2035S, Beyotime, Shanghai, China), followed by DNA-staining with DAPI (4’,6-diamidino-2-phenylindole according to the instruction of this manufacturer (I029-1-1, Nanjing Jiancheng, Nanjing, China). The immunoflorescent staining cells were subjected to confocal microscopic observation of green fluorescent γ-H2AX at λ_ex/em_= 495/519 nm and blue fluorescent DAPI at λ_ex/em_ = 464/454 nm.

### 2.9 Subcutaneous tumor xenografts in nude mice with CDDP intervention

Twenty male nude mice (*BALB/C^nu/nu^* aged at 6 weeks, with 16-18 g in their weights, from the HFK Bioscience, Beijing) were randomly divided into four groups: *WT*_vehicle_, *WT*_CDDP_, *Nrf1α^−/−^*, and *Nrf1α^−/−^*. After normal feeding for 5 d, mouse xenograft models were made by subcutaneous heterotransplantation of HepG2 (*WT*) and *Nrf1α^−/−^* cell lines, respectively, into nude mice as described [31]. Each line of experimental cells (1 × 10^7^) was allowed for its exponential growth and then suspended in 0.2 ml of serum-free DMEM, before being inoculated subcutaneously into the right upper back region of indicated mice at a single site. At the 4^nd^ day inoculated, the indicated groups of mice were subjected to intervention of CDDP (10 mg/kg, by intraperitoneal injection, once). This concentration of chemotherapeutic intervention by CDDP was calculated based on the ratio of human to mouse surface area, as referenced from the treatment guideline of the National Comprehensive Cancer Network (NCCN). Subsequently, the sizes of xenograft tumors were measured every two days and calculated by a standard formula (i.e., V = ab^2^/2) as shown graphically. On the 16^nd^ day after intervention by CDDP, all the mice were euthanized and the tumors were removed, before being subjected to histopathological observations and other experimental analyses. Notably, all the mice were maintained under standard animal housing conditions with a 12-h dark cycle and allowed access *ad libitum* to sterilized water and diet. All relevant studies were carried out on 8-week-old mice (with the license SCXK-PLA-20210211) in accordance with United Kingdom Animal (Scientific Procedures) Act (1986) and the guidelines of the Animal Care and Use Committees of Chongqing University, both of which were also subjected to the local ethical review in China. All relevant experimental protocols were approved by the University Laboratory Animal Welfare and Ethics Committee. Another ethical issue related to the experiment is that the tumor size of a few xenograft model mice, especially in *Nrf1α^−/−^*_vehicle_ group is too large to prevent certain hemorrhagic ulcers and thus shorten the feeding cycle. Overall, such related researches had indeed been conducted in accordance with approved and effective ethical regulations.

### 2.10 Histopathological examination by HE and Tunnel staining

The above-described tumor tissues of each group were fixed with 4% paraformaldehyde, and then transferred to 70% ethanol, according to the conventional protocol. Thereafter, all the individual tumor tissues were placed in the processing cassettes, dehydrated through a serial of alcohol gradient, and then embedded in paraffin wax blocks before being sectioned into a series of 5-µm-thick slides. Next, the tissue sections were dewaxed in xylene, and thereafter washed twice in anhydrous ethanol to eliminate the residual xylene, followed by rehydration in another series of gradient concentrations of ethanol with being distilled. Subsequently, they were stained with the standard hematoxylin and eosin (HE) and visualized by routine microscopy. Furthermore, these tumor tissues were prepared on a slicing machine to yield frozen sections on a glass slide, and then fixed in a constant cold box as described above. Subsequent experimental procedure was manipulated according to the instruction of an *in situ* tunnel fluorescent staining kit (G002-3-1, Nanjing Jiancheng, Nanjing, China), before being stained with DAPI. After the staining solution were removed, these tumor sections were subjected to histopathological observations under a fluorescence microscope.

### 2.11 Oil red staining of lipids

The pre-made sections of tumor tissues were stored in a −20°C refrigerator on a slicing rack. Before experiment, they were allowed to warm up for 5-10 min in room temperature. Thereafter, oil red O staining of these tumor sections was carried out according to the manufacturer’s instructions (D027-1, Nanjing Jiancheng, Nanjing, China) and then subjected to microscopic observation of red lipid droplets.

### 2.12 Bioinformatic analysis of transcriptomes and metabolomes

Total RNAs were extracted from each of experimental cell lines, that had incubated with 3 μM CDDP for 24 h, and then subjected to the transcriptomic sequencing by Beijing Genomics Institute (BGI, Shenzhen, China) on an Illumina HiSeq 2000 sequencing system (Illumina, San Diego, CA). All the detected mRNAs were fragmented into short fragments of approximately 200 bp. The resulting data were bioinformatically analyzed at the BGI database to determine differential expression genes (DEGs) as shown in distinct plotting forms. Thereafter, the untargeted metabolomes were employed to detect distinctions amongst three experimental cell lines (*WT, Nrf1α^−/−^, Nrf2^−/−^*) and between the relevant xenograft tumor tissues (*WT*_vehicle_, *Nrf1α^−/−^*_vehicle_), as conducted in three replicates of each group (Wuhan Metware Biological Co., Ltd). The obtaining data were subjected to bioinformatic analysis of differential metabolites and pathway enrichments. Subsequently, all the results from bioinformatic analyses were further validated by relevant biological experiments

### 2.13 Molecular docking of CDDP within Nrf1α/TCF11

The 3D-structures of indicated ligands were obtained by searching relevant compounds or drugs from the Drugbank (https://go.drugbank.com/drugs/DB00515), whilst the AlphaFold-predicted 3D-structure of Nrf1/TCF11 (of 772 aa, with accession no. Q14494) as a docking receptor macromolecule was searched from the UniPort (https://www.uniprot.org/). The possibly binding pockets of Nrf1/TCF11 were predicted by DeepSite database (https://www.playmolecule.com/deepsite/). These were taken altogether and put into the AUTODOCK Tools1.5.6 software to obtain the ligand-docking information within this macromolecule, which was calculated by a genetic algorithm called Lamarchkian GA (4.2). Subsequently, the docking results are further optimized by both OpenBABEL 3.1.1 and PyMOL softwares.

### 2.14 Statistical analysis

The relevant data presented in this study are shown as fold changes (mean ± SD) relative to indicated controls, that were calculated from at least three and even five independent experiments, each of which was performed in triplicates. Statistical significances were assessed by using one-way ANOVA. Besides Student’s *t*-test, the Tukey’s post hoc test was also employed to determine the statistical significances for all pairwise comparisons of interest. These differences between distinct treatments were considered to be statistically significant at *p* < 0.05 or 0.01.

## 3. Results

### 3.1 Nrf1/TCF11 is predicated to possess a highly binding activity with CDDP

By using a machine-learning procedure, we first simulated and predicted the possibly ligand-binding pockets of Nrf1/TCF11. The resulting data suggest five potential active sites existing in this CNC-bZIP protein, of which two sites with the highest scores were reckoned to perform the molecular docking of CDDP (Figure S1, A & B). That is, the results showed that CDDP could bind to Nrf1 at its amino acid positions of E^403^/Y^712^ (i.e., E^433^/Y^742^ in TCF11) or E^653^ (i.e., E^683^ in TCF11) with their respective binding energy of −7.64 and −7.40, implying a strong binding activity between CDDP and Nrf1/TCF11 (Figure 1B). In addition, it should also be noticed that, in the molecular docking with DTT and tBHQ (Figure S1, C & D), their binding energy to Nrf1/TCF11 was estimated to be only a half of the CDDP’s binding energy to the CNC-bZIP protein. In fact, it had been proved in previous studies on DTT and tBHQ [22, 23], that both compounds can indeed affect the regulation of those redox signaling networks mediated by Nrf1. Next, we continue to further experimentally validate the virtual impact of CDDP on the expression levels of Nrf1 and its transcriptional activity to regulate its cognate target genes involved in diverse biological processes during development and even malignant progression of liver cancer.

### 3.2 Distinct induction of Nrf1 and Nrf2 by CDDP, that inhibits the proliferation of different cell lines

As shown in Figure 1C, CDDP exerted significant inhibitory effects on the cell viability and even proliferation of *WT, Nrf1α^−/−^* and *Nrf2^−/−^* lines, which were dependent on the concentrations of CDDP and its intervention times. Such distinct genotypic cell lines exhibited different responses to CDDP in the same conditions. Amongst them, *WT* cells reached the *IC_50_* of CDDP at a concentration of 15.6 μM, but loss of *Nrf1α* or *Nrf2* did not appear to cause a significant increase in such induced death of *Nrf1α^−/−^* and *Nrf2^−/−^* cell lines that had been treated with 0.75 μM to 25 μM of CDDP for 24 h (Figure 1C, *c1*), albeit their deaths gradually incremented and became slightly significant after the doses of CDDP were increased from 50 μM to 100 μM. Thereby, the *IC_50_* concentrations of CDDP for 24-h intervention of both *Nrf1α^−/−^* and *Nrf2^−/−^* cell lines were evaluated to be much higher than 50 μM. When the CDDP intervention time was extended to 48 h, its inhibitory effects on these cell lines become stronger, especially after being treated from 1.5 μM to 25 μM of this drug, such that growth of these cell lines was sharply decreased to a certain maximal extent (Figure 1C, *c2*), but then become almost unaffected by increasing doses of CDDP to 50∼100 μM. In addition, almost no marked differences in the response of these three cell lines to 48-h intervened CDDP were observed, relative to those observed after its 24-h treatment.

Through validation of these three cell lines by Western blotting as described previously [6], it was confirmed truly that *Nrf1α^−/−^* cells gave rise to a loss of Nrf1α protein expression, but along with a significant increase in the expression of Nrf2 when compared with that measured in *WT* cells (Figure 1D). The loss of Nrf2 in the resulting *Nrf2^−/−^* cells led to relatively weakened expression of Nrf1α, as compared to *WT* controls. Here, it was also found that distinct inducible expression patterns of Nrf1 and Nrf2 in CDDP-intervened *WT* cells depended on different doses of this compound and its treatment time courses (Figure 1E). Of sharp note, strikingly increased expression of Nrf1α proteins occurred primarily after treatment of *WT* cells with 1.5∼6 μM of CDDP for 24 h (Figure 1E, *e1*) or 3 μM of CDDP for 4 h to 12 h (*e4*). By contrast, only modest induction of Nrf2 appeared to be stimulated by 1.5 μM of CDDP, but its expression was then gradually decreased or almost disappeared after the doses of this drug were increased to 6 μM (Figure 1E, *e2*). As such being the case, the time course of 3 μM CDDP-stimulated Nrf2 expression was roughly similar to the course of Nrf1α induction by this compound (Figure 1E*, c.f. e5 with e4*).

Next, to examine distinct genotypic cellular responses to 3 μM CDDP intervention for 12 h or 24 h, it was unraveled d that the protein expression levels of Nrf1α were significantly increased after 12-h treatment of *WT* cells, and then modestly decreased at 24 h of this treatment, to some certain degrees that were still higher than those measured at *T_0_* (Figure 1F, *upper panel*). The transcriptional expression of *Nrf1* was markedly induced by CDDP intervention for 12 h or 24 h, with an exception for being modestly decreased by its longer intervention for 24 h (Figure 1G), whereas the transcriptional expression of *Nrf2* was only marginally induced by CDDP treatment in *WT* cells. Although the overall abundances of Nrf1α in *Nrf2^−/−^* cells were considerably lowered when compared with its expression levels in *WT* cells, its remaining protein expression was still slightly induced by CDDP (Figure 1F, *upper panel*). Besides, the mRNA expression levels of *Nrf1* were also modestly stimulated by this intervention of *Nrf2^−/−^* cells (Figure 1G). However, knockout of *Nrf1α^−/−^* led to the aberrant overexpression of Nrf2 at its protein levels (Figure 1F, *middle panel*) and mRNA levels (Figure 1G), and such highly-expressed abundances of Nrf2 were significant reduced by CDDP, relative to their *T_0_* control values.

Further intervention of *Nrf2^−/−^* cells with different doses of CDDP for 24 h caused an inducible increase in the Nrf1α expression, stimulated obviously by 1.5 μM of CDDP, and then its induced expression levels were gradually decreased with increasing doses of this drug (Figure 1H, *h1*). The time course of 3 μM CDDP-chasing experiments revealed that the protein expression levels of Nrf1 were gradually increased from 4 h to 12 h of this intervention, and then gradually decreased until 24 h of the experiments stopped, albeit its abundances were much fainter in *Nrf2^−/−^* cells as compared to the *WT* value measured at *T_4h_* (Figure 1H, *h4*). Conversely, further examination of the Nrf2 protein expression in CDDP-intervened *Nrf1α^−/−^* cells uncovered a dose-dependent and also time-dependent inhibitory effect of this drug on the CNC-bZIP expression (Figure 1I, *i1 & i4*). Of note, intervention of 3 μM CDDP led to a biphasic expression pattern of Nrf2, which was first suppressed by this treatment for 4 h to 8 h, and then gradually recovered to a considerably higher extent, closer to its *T_0_* level (Figure 1I, *i4*).

Further experiments revealed distinct stimulated changes in the expression of those redox genes (regulated by Nrf1 and/or Nrf2) in distinct cellular responses to CDDP (Figure 1, J & K). Of note, both the protein and mRNA expression levels of HO-1 were induced by CDDP intervention of *WT* cells for 12 to 24 h. The loss of *Nrf1α* led to an obvious increase in the basal protein and mRNA abundances of HO-1 in *Nrf1α^−/−^* cells, and its further inducible increased expression occurred only after CDDP intervention for 12 h, but then such CDDP-stimulated expression was suppressed by 24-h treatment with this drug to a considerably lower level, close to the *T_0_* control of *WT* cells. Conversely, CDDP-inducible protein and mRNA expression of HO-1 appeared to be abolished or further repressed in *Nrf2^−/−^* cells (Figure 1, J & K). By contrast, transcriptional expression of *GCLM* and *GSR,* but not *GPX1,* was also up-regulated by CDDP intervention of *WT* cells for 12 h to 24 h, whilst both *NQO1* and *GCLC* were significantly down-regulated by this drug (Figure K). Although such these genes (i.e., *NQO1, GCLC, GCLM, GPX1* and *GSR*) were basally up-regulated in *Nrf1α^−/−^* cells, they appeared to be suppressed or unaffected by CDDP treatment. In turn, basal mRNA expression of *NQO1, GCLC, GPX1, GSR,* but not *GCLM*, was significantly down-regulated in *Nrf2^−/−^* cells, but they were allowed for differential responses to CDDP intervention; *GCLC* and *GCLM* were apparently induced, while *GSR* and *NQO1* were further down-regulated, along with *GPX1* being unaffected, by this drug. As such being the case, similar but different protein expression patterns of these redox genes regulated by Nrf1 and Nrf2 in response to CDDP were also demonstrated (as shown in Figure 1J). Taken together, it is inferable that Nrf1 and Nrf2 make their unique and coordinated contributions to the antioxidant and detoxifying responses to CDDP, but the inducible transactivation of Nrf1 is likely to play a key role in this process (as deciphered in a proposed model, see Figure 1L).

### 3.3 Discrete responses of Nrf1 and Nrf2 to genotoxic and cytotoxic effects of CDDP in different cell lines

As shown in Figure 1L, it is clearly known that genotoxic and cytotoxic effects of CDDP are triggered primarily by this drug-caused DNA damage and redox imbalance [29, 32, 33]. Hence, we examined changes in the DNA damage to different genotypic (*WT, Nrf1α^−/−^*, *Nrf2^−/−^*) cell lines, after they had been intervened by CDDP for 24 h. The results by immunofluorescent staining of γ-H2AX (as a typical marker of the nuclear DNA damages) revealed that CDDP led to an obvious enhancement of the DNA damage in *Nrf2^−/−^* cells (in which Nrf1 was markedly down-regulated), when compared with that of *WT* cells, although the *WT* cellular DNA damage was also modestly enhanced by this drug, relative to its vehicle control (Figure 2A). As such, CDDP-induced DNA damage seemed to be, however, mitigated to a certain extent by loss of Nrf1α (in *Nrf1α^−/−^* cells retaining aberrant accumulation of the hyperactive Nrf2). Largely similar results were obtained from Western blotting of total H2AX and its phosphorylated γ-H2AX, revealing significant decreases in their basal and CDDP-inducible expression abundances in *Nrf1α^−/−^* cells, when compared with those levels of such proteins expressed in *WT and Nrf2^−/−^* cell lines (Figure 2B, *b2* & *b3*). Of note, the CDDP-induced γ-H2AX, but not its basal, levels were also marginally lowered in *Nrf2^−/−^* cells, relative to those counterparts of *WT* cells. Furthermore, the real-time qPCR data unraveled loss of *Nrf1α* caused the basal, but not CDDP-stimulated, mRNA expression of *H2AX* to be down-regulated in *Nrf1α^−/−^* cells, but both mRNA expression levels of *H2AX* were significantly up-regulated in *Nrf2^−/−^* cells when compared to *WT* cells (Figure 2B, *upper panel*). Collectively, these indicate that Nrf1, rather than Nrf2, is required for governing the stable expression of *H2AX*, albeit both factors are essentially involved in the cytoprotective response to the DNA damage caused by CDDP.

**Figure 2.**
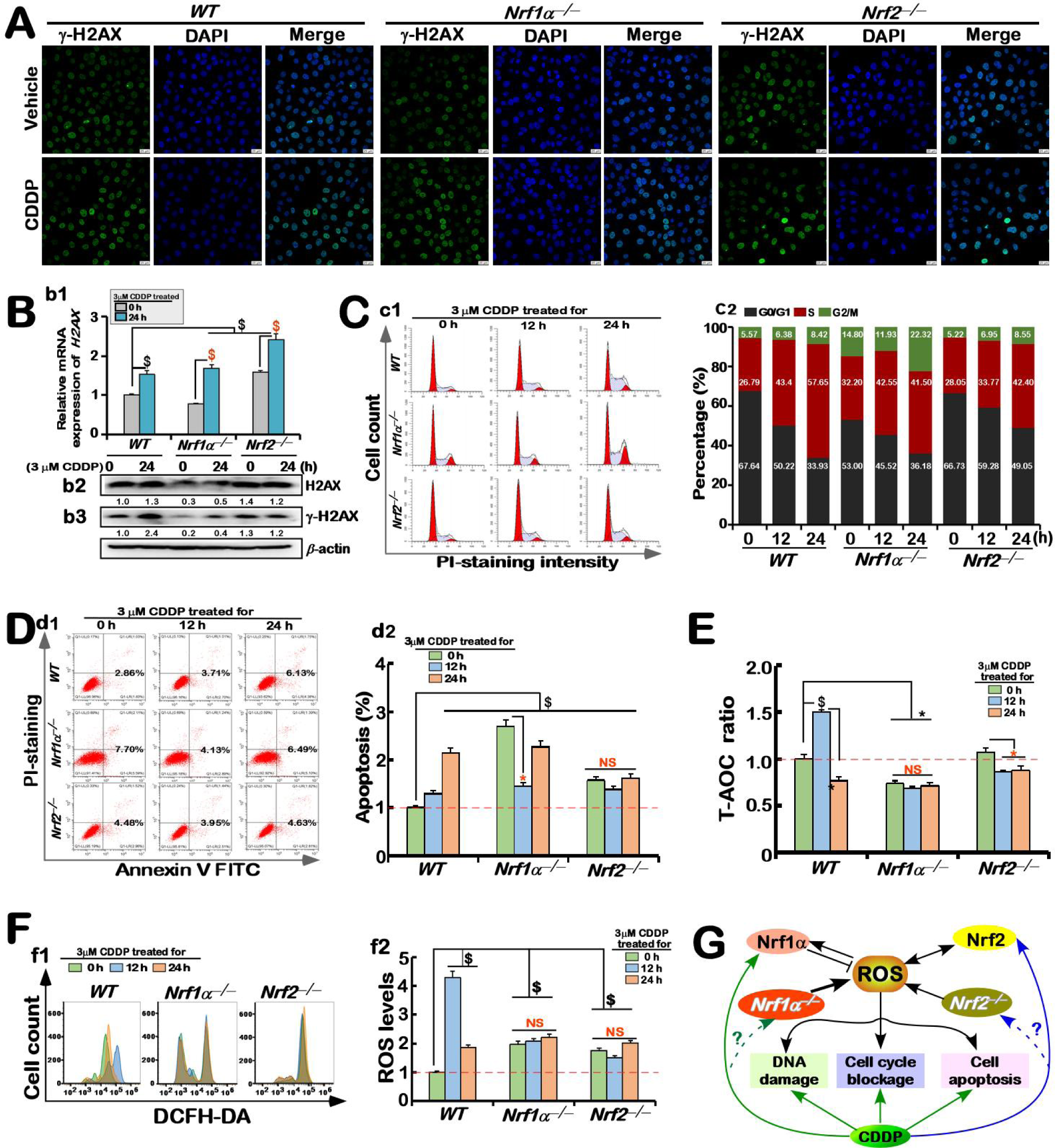
Distinct genotypic cellular responses of Nrf1 and Nrf2 to genotoxic and cytotoxic effects of CDDP. **(A)** Immunofluorescent staining of γ-H2AX that was used to detect the DNA damage of different genotypic (*WT, Nrf1α^−/−^* and *Nrf2^−/−^*) cell lines, which was caused by CDDP or vehicle (i.e., Complete medium with solvent). Subsequently, the DNA staining was conducted with DAPI. **(B)** Basal and CDDP-stimulated expression of H2AX at mRNA **(*b1*)** and protein **(*b2*, *b3*)** levels were determined by real-time qPCR and Western blotting with distinct antibodies against H2AX or γ-H2AX (i.e., phosphorylated H2AX, which serves as a typical DNA damage marker), respectively. β-actin served as a loading control. The intensity of these immunoblots was quantified and the real-time qPCR data were statistically analyzed, as described in the legend for Figure 1. **(C)** Distinct effects of CDDP on different cell cycles were evaluated by flow cytometry, after these cell lines (*WT, Nrf1α^−/−^* and *Nrf2^−/−^*) had been intervened with 3 μM of CDDP for 0, 12, or 24 h. The resulting data are shown in the column charts (***c2***), which were representative of at least three independent experiments being each performed in triplicate. **(D)** Distinct apoptotic effects of CDDP on different cell lines (*WT, Nrf1α^−/−^* and *Nrf2^−/−^*) were also determined by flow cytometry, after these cell lines had been treated with 3 μM CDDP for different time periods (*i.e.*, 0, 12 and 24 h). Subsequent experimental procedure was carried out as described in the “Method and materials”. The resulting data were analyzed and also shown in the column charts **(*d2*)**, which were representative of at least three independent experiments being each performed in triplicate. **(E)** CDDP-caused changes in the total antioxidant capacity (T-AOC) were determined in three distinct cell lines that had been treated with 3 μM of CDDP for 0, 12 or 24 h. The resulting absorbance values (*OD*) were tested, and then the T-AOC ratio was calculated as shown graphically, along with statistical significances being analyzed. **(F)** Flow cytometry analysis of intracellular ROS levels in *WT*, *Nrf1α^−/−^* and *Nrf2^−/−^* cell lines, that had been treated with 3 μM CDDP for 0, 12 or 24 h, by determining the DCFH-DA fluorescent intensity (***f1***). The resulting data were further analyzed by the FlowJo 7.6.1 software as shown graphical (***f2***), with significant increases ($$, *p* < 0.01 and $, *p* < 0.05), significant decreases (* *p* < 0.05, \*\**p* < 0.01) and no significant differences (NS), which were statistically determined by comparing each basal value of [*Nrf1α^−/−^*] and [*Nrf2^−/−^*] with that of *WT*t0. Symbols ‘$ or *’ also indicate significant differences in *Nrf1α^−/−^* or *Nrf2^−/−^* cells, when compared to their respective *t0* after CDDP intervention. **(G)** A schematic representation of distinct responses of Nrf1 and Nrf2 to the ROS products resulting from CDDP, in order to maintain intracellular redox homeostasis. Since the loss of *Nrf1α^−/−^* or *Nrf2^−/−^* results in certainly varying degrees of oxidative stress, this exacerbates the CDDP-caused DNA damage, cell cycle arrest, and apoptosis. CDDP acts as a functional inducer for DNA damage, allowing cancer cells to death, but inconsistent effects of *Nrf1α^−/−^*or *Nrf2^−/−^* in this process are worthwhile for further studies.

Before CDDP intervention, flow cytometry analysis showed a significantly lowered proportion (53.0%) of the G0/G1 phase in *Nrf1α^−/−^* cells, when compared to equivalents of both *WT* (67.64%) and *Nrf2^−/−^* (66.73%) cell lines (Figure 2C, *c1 & c2*). Conversely, the loss of *Nrf1α* gave rise to two higher proportions in the S (32.2%) and G2/M (14.8%) phases of *Nrf1α^−/−^*cells than those counterparts of *WT* (26.79%, 5.57%) and *Nrf2^−/−^* (28.05%, 5.22%). This indicates that *Nrf1α^−/−^*cells are required for certain longer times of its DNA, RNA and protein synthesis in order to meet the needs of its enhanced cell division and malignant proliferation, which is likely explained as a reason for leading to abnormal nutritional metabolism activities of *Nrf1α^−/−^* cells [34]. Next, the impact of cytotoxic CDDP on these cell cycles was examined herein. The results revealed that CDDP intervention for 24 h caused a substantial inhibitory effect on the *WT*’s G0/G1 phase to 33.9%, whereas the G0/G1 phases of *Nrf1α^−/−^*or *Nrf2^−/−^* cell lines were only partially retarded by this drug to 36.18 % or 49.05%, respectively (Figure 2C, *right panel*). By contrast, the *WT’s* S-phase was significantly arrested by CDDP, allowing its proportion to be extended to 57.65%, whilst the S-phases of *Nrf1α^−/−^* or *Nrf2^−/−^* cells were only modestly retarded by this drug to 41.5% or 42.4% proportioned. Of note, the G2/M phase of *Nrf1α^−/−^* cell cycle was significantly augmented to 22.32% by CDDP intervention for 24 h, but no obvious changes in the G2/M phases of *WT* and *Nrf2^−/−^* cell lines were shown (in Figure 2C, *c2*). Overall, these demonstrate that the G2/M phase of *Nrf1α^−/−^*, but not *Nrf2^−/−^*, cells is arrested by CDDP to meet the needs of increasing mitotic division cycles, even albeit its S-phase was partially shortened to hinder its growth, whereas both *WT* and *Nrf2^−/−^* cell cycles are strikingly arrested by this drug at their S and/or G0/G1 phases insomuch as to slow down their growth.

The effect of cytotoxic CDDP on apoptosis of distinct cell lines was also examined by flow cytometry. It was found that apoptosis of *WT* cells was increased by CDDP intervention for 12 h to 24 h (Figure 2D, *d1* & *d2*), but CDDP-stimulated apoptosis of *Nrf1α^−/−^* cells was biphasic (which was down-regulated by 12-h intervention with this drug and then mostly recovered by its extended intervention for 24h), whilst *Nrf2^−/−^* cell apoptosis was almost unaffected by this drug, albeit basal apoptosis of *Nrf1α^−/−^*and *Nrf2^−/−^* cell lines was enhanced relative to that of *WT* cells (Figure 2D, *d2*). These are roughly consistent with the aforementioned results of cell proliferation activity in response to CDDP, indicating that the cytotoxicity of CDDP appears to be certainly alleviated and even abolished by loss of Nrf1α and/or Nrf2. Particularly in *Nrf2^−/−^* cells, the remnant Nrf1 factor was induced by CDDP and then exerted its robust cytoprotective role in defending against cytotoxic and genotoxic effects of this drug, while apoptosis of *Nrf1α^−/−^* cells were only temporarily suppressed by hyperactive Nrf2 stimulation by CDDP for 12 h and then partially recovered for 24 h in a time-dependent manner.

Further examination uncovered such a marked reduction in basal level of *t*otal *<u>a</u>*nti*<u>o</u>*xidant *<u>c</u>*apacity (T-AOC) in *Nrf1α^−/−^* cells when compared to that of *WT* or *Nrf2^−/−^* cell lines, and the reduced T-AOC was almost unaffected by CDDP treatment for 12 to 24 h (Figure 2E). However, CDDP-enhanced T-AOC was determined only after its intervention of *WT* cells for 12 h and then repressed steeply to a considerably low extent than its basal ROS levels. An additional inhibitory effect of CDDP on the T-AOC were also determined in *Nrf2^−/−^* cells (Figure 2E). Collectively, these indicate that both Nrf1 and Nrf2 are required for mediating antioxidant responses to CDDP. Accordingly, basal levels of intracellular ROS (as determined by flow cytometry analysis of DCFH-DA) were obviously elevated in *Nrf1α^−/−^* or *Nrf2^−/−^* cell lines, but such higher ROS levels were largely unaffected by CDDP treatment for 12∼24 h (Figure 2F, *f1 & f2*). By contrast, CDDP-inducible ROS levels were increased only after 12-h treatment of *WT* cells and thereafter abruptly reduced by its prolonged intervention for 24 h to considerably lowered extents (albeit being still higher than its basal levels). Taken altogether, these results demonstrate that both Nrf1 and Nrf2 can mediate effective antioxidant, detoxification and cytoprotective responses to the genotoxic and cytotoxic effects of CDDP, because this drug is enabled to cause temporary increases in the intracellular ROS and concomitantly T-AOC of *WT* cells.

### 3.4 Diverse induction of redox-related genes regulated by Nrf1 or Nrf2 in different cellular responses to CDDP

From the above-described results, it is worth questioning why such deficiencies of *Nrf1α* and/or *Nrf2* cannot exacerbate, but conversely resist or even ameliorate, the cytotoxic effects of CDDP on *Nrf1α^−/−^* or *Nrf2^−/−^* cell lines (Figure 1G), even though this drug caused a significant decrease in the viability of *WT* cells, with time-dependent changes in the cell apoptosis, cell cycle, T-AOC and ROS levels. This also raises a doubt about why their responses to CDDP appeared to be not consistent with previous studies [35], revealing that severe oxidative stress remains to take place in *Nrf1α^−/−^* cells, where Nrf2 was accumulated and those antioxidant genes were also concomitantly up-regulated, however. To address this, we thereby continued to investigate such discretely-stimulated effects of CDDP intervention on the expression of those redox-related genes regulated by Nrf1 or Nrf2 in distinct genotypic cell lines.

The transcriptome sequencing data revealed significantly CDDP-stimulated changes in differential expression genes responsible for redox regulation in *WT* cells, when compared to those of untreated cells (Figure 3A), while there appeared to be relatively less changes in such redox gene expression in CDDP-intervened *Nrf1α^−/−^* or *Nrf2^−/−^* cell lines. Subsequently, real-time qPCR data further validated that CDDP acted as a particularly effective inducer to trigger the significant mRNA expression of those redox-regulatory genes in *WT* cells (Figure 3, B-F). Such genes included those responsible for ROS scavengers (e.g., *CAT, PRDX1, PRDX3, PGX2, SOD1, SOD2* in Figure 3B), NADPH biosynthesis (e.g., *IDH1, IDH2, ME1, TIGAR* in Figure 3C), GSH biosynthesis and transport (e.g., *GCLC, GCLM, GLS,* except for *SLC7A11* being down-regulated in Figure 3D), thioredoxin antioxidant system (e.g., *TXN1, TXN2, SRXN1* in Figure 3E) and the others involved in the iron-based redox homeostasis (e.g., *HO1, SESN1, SESN3* in Figure 3F).

**Figure 3.**
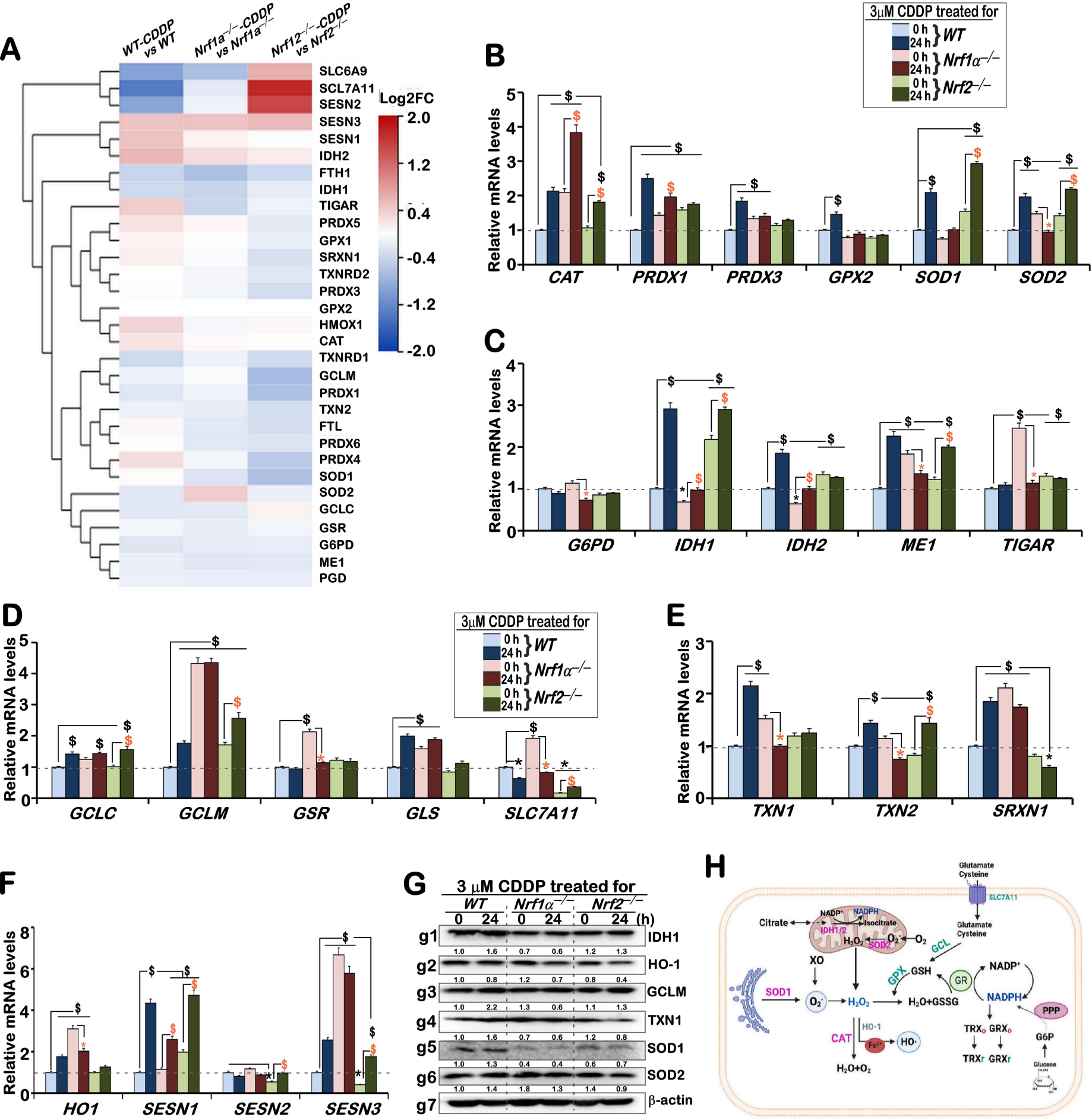
Discrete induction of Nrf1/2-mediated redox-related gene expression in different cellular responses to CDDP. **(A)** The heat-map shows differential expression levels of those antioxidant and detoxification genes determined by transcriptome sequencing of *WT*, *Nrf1α^−/−^* and *Nrf2^−/−^* cell lines that had been intervened with 3 μM CDDP for 0 or 24 h. **(B-F)** Real-time qPCR analysis of basal and CDDP-stimulated expression levels of those genes responsible for redox homeostasis, e.g., (B) balancing the ROS production and elimination (*CAT, PRDX1, PRDX3, GPX2, SOD1, SOD2*), (*C*) NADPH synthesis genes (*G6PD, IDH1, IDH2, ME1, TIGAR*), **(*D*)** GSH biosynthesis and transport (*GCLC, GCLM, GSR, GLS, SLC7A11*), **(*E*)** thioredoxin (TXN) antioxidant system (*TXN1, TXN2, SRXN1*) and **(*F*)** Iron homeostasis (*HO-1, SESN1, SESN2, SESN3*) in *WT*, *Nrf1α^−/−^* and *Nrf2^−/−^* cell lines that hat had been intervened with 3 μM CDDP for 24 h or not. The resulting data were shown as fold changes (mean ± SD, n = 3 × 3), which are representative of at least three independent experiments being each performed in triplicates, with significant increases ($$, *p* < 0.01 and $, *p* < 0.05), significant decreases (* *p* < 0.05, \*\**p* < 0.01) and no significant differences (NS), as compared each basal value of [*Nrf1α^−/−^*] and [*Nrf2^−/−^*] with that of *WT*t0. Symbols ‘$ or *’ also indicate significant differences in *Nrf1α^−/−^*or *Nrf2^−/−^* cells, when compared to their respective *t0* after CDDP intervention. **(G)** Western blotting analysis of the indicated proteins, including IDH1, HO-1, GCLM, TXN1, SOD1 and SOD2 in *WT*, *Nrf1α^−/−^*and *Nrf2^−/−^* cell lines that had been intervened with 3 μM CDDP for 0 or 24 h. The intensity of these immunoblots was also quantified by the Quantity One 4.5.2 software and showed under those indicated protein bands. β-actin served as a loading control. **(H)** A schematic representation of redox signaling pathway to balance intracellular production and elimination of ROS, including O2·, H2O2, O2^−^, and OH·.

Albeit hyperactive Nrf2 was accumulated in *Nrf1α^−/−^*cells, its basal and CDDP-inducible up-regulation of only *CAT*, *PRDX1, SESN1* were determined (Figure 1, B & F). But, stimulated expression of *SOD2*, *ME1, TIGAR, SLC7A11, GSR, TXN1, TXN2, HO-1* and *SESN3* was significantly suppressed by CDDP in *Nrf1α^−/−^* cells, even though their basal expression levels were also up-regulated to certain extents (Figure 1, B to F). This indicates these genes are differentially regulated by Nrf2, most of which are concomitantly down-regulated by CDDP-inhibited expression of Nrf2. Conversely, CDDP-intervened *Nrf2^−/−^* cells gave rise to significant up-regulation of *CAT*, *SOD1, SOD2, IDH1, ME1, GCLC, GCLM, TXN2, SESN1* and *SESN3* (Figure 1, B to F), which is likely attributable to the remaining Nrf1 stimulation by this drug.

Further Western blotting revealed that basal abundances of HO1, GCLM, TXN1 and SOD2 were up-expressed in *Nrf1α^−/−^*, when compared to their counterparts of *WT* and *Nrf2^−/−^* cell lines (Figure 3G), but CDDP intervention caused significant down-regulation of HO-1 and SOD2 in *Nrf1α^−/−^* cells, whilst HO-1, TXN1 and SOD2 were also down-regulated by this drug in *Nrf2^−/−^* cells. Conversely, most of these examined proteins, e.g., IDH1, GCLM, TXN1, SOD1 and SOD2 were up-regulated in CDDP-treated *WT* cells (Figure 3G). Of note, both basal and CDDP-inducible expression levels of SOD1 were almost abolished by the loss of *Nrf1α^−/−^* or *Nrf2^−/−^* (Figure 3G, *g5*). Taken together, these indicate that Nrf1 and Nrf2 are differentially contributable to redox gene regulation in response to CDDP.

### 3.5 Distinctive requirements of Nrf1 and Nrf2 for different cellular signaling responses to CDDP

As shown in Figure 4A, CDDP-stimulated expression levels of those critical genes required for multiple signaling pathways in the cellular response to its genotoxicity and cytotoxicity were analyzed by transcriptome sequencing. Amongst them, five genes *FGF21, DDIT3, SNAI2, JUN* and *FOS* were significantly up-regulated by CDDP in *Nrf2^−/−^*cells, whereas only MMP9 was modestly up-regulated by this compound in *Nrf1α^−/−^*cells, but down-regulated in the intervened *Nrf2^−/−^* cells (Figure 4A). Further real-time qPCR analysis unraveled that *Nrf1α^−/−^* cells gave rise to marked decreases in basal expression levels of most examined genes involved in unfold protein responses (UPRs), such as *PDI, BIP, PERK, EIF2A, ATF4, XBP1*, *ATF6, CHOP, FGF21* and *HSP60*, of which the others, except for *ATF6*, were also basally down-regulated by *Nrf2^−/−^*cells, when compared to their counterparts of *WT* cells (Figure 4B). Upon CDDP intervention, its inducible mRNA expression of *IRE1*, *CHOP* and *FGF21* was significantly augmented in this drug-treated *WT* or *Nrf2^−/−^*cells for 12 h, but then abruptly down-regulated by its prolonged intervention for 24 h to considerably lower levels than their respectively basal levels, while *ATF6* was only marginally induced by CDDP intervention of *WT* or *Nrf2^−/−^* cells for 12 to 24 h (Figure 4B). By contrast, the expression of *IRE1* was basally up-regulated, but substantially down-regulated by CDDP in *Nrf1α^−/−^* cells. In addition, CDDP-inducible expression levels of the other genes in *Nrf1α^−/−^* or *Nrf2^−/−^* cells remained hardly close to their corresponding basal levels of *WT* cells.

**Figure 4.**
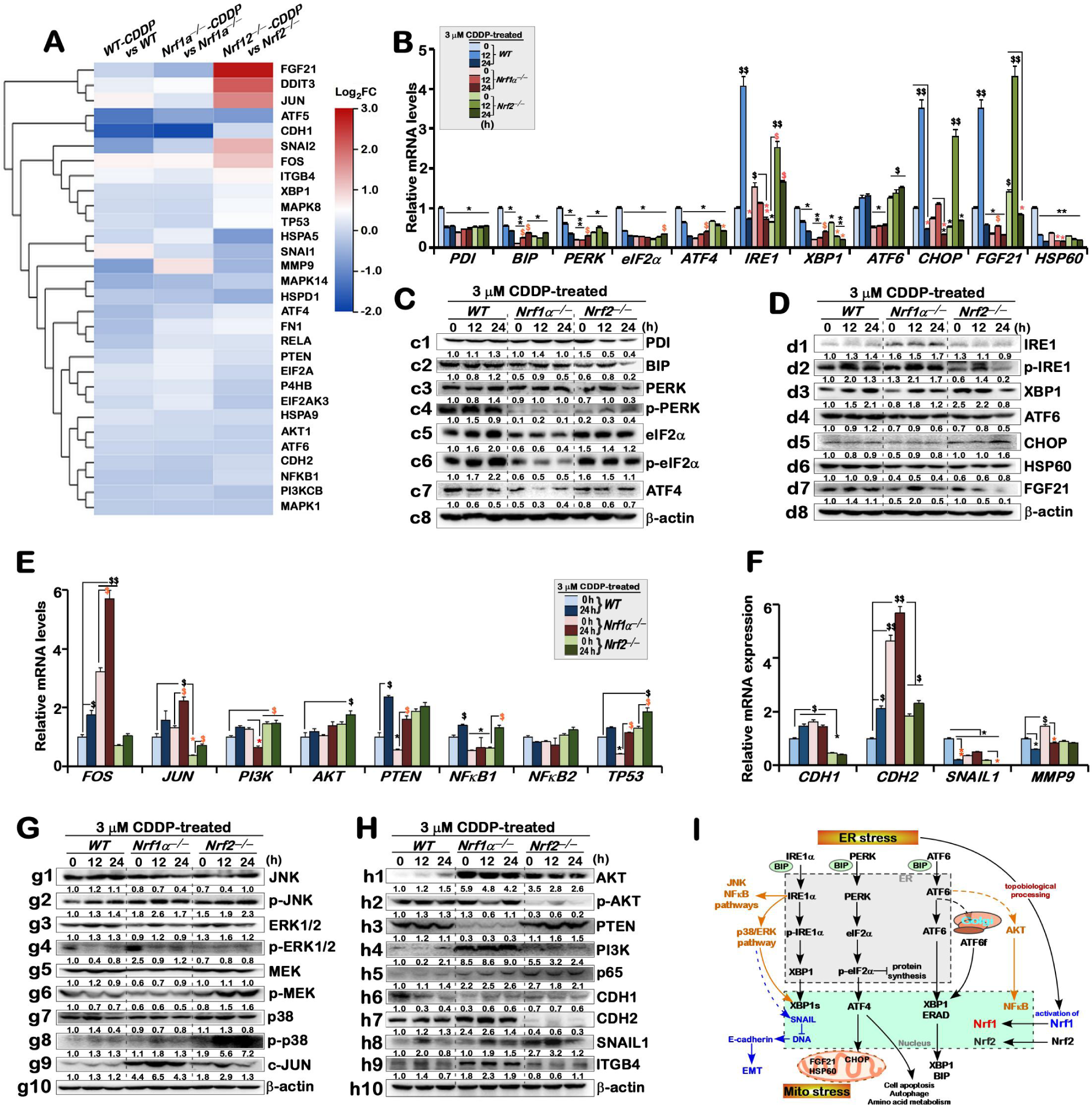
Distinctive requirements of Nrf1 and Nrf2 for different cellular signaling responses to CDDP. **(A)** The heat-map shows changes of those differential expression genes required for multi-hierarchical signaling networks (e.g., UPR, MAPK, PI3K/AKT-PTEN and EMT pathways), which were determined by transcriptome sequencing of *WT*, *Nrf1α^−/−^*and *Nrf2^−/−^* cell lines that had been intervened with 3 μM CDDP. **(B)** Real-time qPCR analysis of the mRNA expression levels of those genes responsible for the endoplasmic reticulum (ER) and/or mitochondrial stresses (*PDI, BIP, PERK, eIF2α, ATF4, IRE1α, XBP1, ATF6, CHOP, HSP60* and *FGF21*) in *WT*, *Nrf1α^−/−^* and *Nrf2^−/−^* cell lines that had been treated with 3 uM CDDP for 0, 12 or 24 h. **(C,D)** Western blotting analysis of those expression abundances of indicated proteins involved in the ER and mitochondrial stress responses after CDDP stimulation of *WT*, *Nrf1α^−/−^* and *Nrf2^−/−^* cell lines for 0, 12 and 24 h. β-actin served as a loading control. **(E,F)** Real-time qPCR analysis of basal and CDDP-stimulated expression levels of those genes responsible for MAPK, PTEN-PI3K-AKT (E) and EMT (F) signaling pathways in *WT*, *Nrf1α^−/−^* and *Nrf2^−/−^* cell lines that had been intervened by this drug for 0 or 24 h. **(G,H)** Abundances of those indicated proteins involved in the MAPK, PTEN-PI3K-AKT and EMT signaling pathways were determined by Western blotting of *WT*, *Nrf1α^−/−^* and *Nrf2^−/−^* cell lines that had been intervened by this drug for 0, 12 or 24 h. β-actin served as a loading control. (I) A schematic diagram of multiple signaling pathways in the putative cellular responses to stress stimulated by CDDP. They include the unfold protein response (UPR) to the ER and mitochondrial stresses, the PTEN-PI3K-AKT signaling cascades, those signaling pathways mediated by NF-κB and MAPKs (i.e., JNK, ERKs, and p38), and the epithelial to mesenchymal transition (EMT)-related signaling. All these signaling transduction pathways were referenced from the KEGG pathway and cross-talked together to form multi-hierarchical signaling networks. Besides Nrf2, Nrf1 is also likely processed to be activated and then translocated into the nucleus before regulating its target genes. Note: The intensity of those immunoblots was quantified by the Quantity One 4.5.2 software and also showed below the indicated protein bands. The RT-qPCR resulting data were shown as fold changes (mean ± SD, n = 3 × 3), which are representative of at least three independent experiments being each performed in triplicates. Significant increases ($, p < 0.05; $$, p < 0.01) and significant decreases (\**p* < 0.05; \*\**p* < 0.01) were statistically determined, when compared with *WT* controls (measured at 0 h), respectively. Additional “$ or *” symbols indicate significant differences in *Nrf1α^−/−^* or *Nrf2^−/−^* cell lines that had been intervened by CDDP as compared with their respective *t0* values after intervention.

Western blotting analysis of the above-described gene products revealed that CDDP intervention of *WT* cells caused obvious increases in abundances of PDI (also called P4HB, a dithioredoxin-containing isomerase inhibiting misfolded protein aggregation [23]), BIP (known as GRP78, a key player in the protein folding and quality control of the ER), PERK, elF2α, IRE1, XBP1, ATF6, FGF21, as well as three phosphorylated p-PERK, p-elF2α and p-IRE1, to differentially varying extents (Figure 4, C & D). In *Nrf1α^−/−^* cells, only both p-IRE1 and XBP1 were significantly up-regulated by CDDP treatment for 12 to 24 h, apart from PDI, BIP and FGF21 being up-regulated only after this treatment for 12 h, but not for 24 h. However, basally down-regulated abundances of p-PERK, elF2α, p-elF2α, ATF4, CHOP and HSP60 (a player in the correct assembly of misfolded or unfolded proteins under mitochondrial stress) were detected in *Nrf1α^−/−^* cells (Figure 4C &D), and they appeared to be unaffected or further suppressed by CDDP, while basally up-regulated IRE1 expression was also unaffected by this drug (Figure 4D). In *Nrf2^−/−^* cells, induction of PERK, p-PERK, p-IRE1 and CHOP by CDDP occurred primarily after this intervention for 12 h, rather than 24 h except for CHOP, whereas PDI and FGF21 were apparently diminished by this intervention, aside from that basally enhanced elF2α and p-elF2α were almost unaltered by this drug.

Further experiments by real-time qPCR uncovered significant increases in the mRNA expression levels of *FOS, PTEN* and *CDH2* was stimulated by CDDP in *WT* cells, whilst *JUN, PI3K (p110), NFκB1, TP53 and CDH1* were only modestly up-regulated, but *SNAIL1* and *MMP9* were down-regulated by this intervention (Figure 1, E & F). By contrast, both basal and CDDP-inducible levels of *FOS, JUN,* and *CDH2* were further augmented in *Nrf1α^−/−^* cells, and the basally lowered *PTEN* and *TP53* were also partially induced by CDDP, but the basally elevated *PI3K (p110)* and *MMP9* were suppressed by this drug (Figure 1, E & F). In *Nrf2^−/−^* cells, *AKT*, *NFκB1* and *TP53* were modestly up-regulated by CDDP, whereas *PI3K (p110)* and *PTEN* were unaffected, albeit the latter two expression was basally enhanced to certain higher extents. Basal and inducible levels of *SNAIL1* and *CDH1* were almost abolished by the loss of *Nrf2^−/−^*.

Next, to investigate putative impacts of Nrf1 and Nrf2 on the MAPK signaling responses to CDDP, the relevant protein expression levels were determined by Western blotting (Figure 4G). The abundances of JNK, p-JNK, MEK, ERK1/2, and p38 kinase were induced by CDDP, primarily after its intervention of *WT* cells for 12 h. The induction of p-JNK by CDDP was detected, though total JNK was unaffected or reduced by this drug, in *Nrf1α^−/−^* or *Nrf2^−/−^*cell lines (Figure 4G, *cf. g2 with g1*). The total abundances of ERK1/2 were modestly up-regulated by CDDP intervention of *Nrf1α^−/−^* or *Nrf2^−/−^*cell lines for 12 h, but its phosphorylated protein p-ERK1/2 levels were almost completely abolished by this intervention of all three cell lines (Figure 4G, *cf. g4 with g3*), no matter whether basal p-ERK1/2 levels were enhanced in *Nrf1α^−/−^* cells or reduced in *Nrf2^−/−^* cells, respectively. Also, both MEK and p-MEK was basally down-regulated in *Nrf1α^−/−^* cells and unaffected by CDDP (Figure 4G, *g6 vs g5*), while p-MEK and p-38 kinase were only significantly induced by this drug in *Nrf2^−/−^* cells (*g6, g8*). However, both basal and CDDP-stimulated expression levels of c-JUN (part of AP1, as a target gene involved in tumorigenesis) were significantly augmented in *Nrf1α^−/−^* cells, when compared to its equivalents of *WT* and *Nrf2^−/−^* cell lines (Figure 4G, *g9*).

By immunoblotting analysis of the PTEN-PI3K-AKT signaling response to CDDP, it was found that the protein expression levels of AKT and PI3K (p110) were significantly increased by the loss of *Nrf1α^−/−^*, while the expression of tumor suppressor PTEN was significantly diminished in *Nrf1α^−/−^* cells (Figure 4H, *h1, h3, h4*). Upon intervention with CDDP, the AKT expression was modestly reduced in *Nrf1α^−/−^* or *Nrf2^−/−^*cells, but conversely increased in *WT* cells. By contrast, the phosphorylated protein p-AKT was also modestly induced by CDDP in *WT* cells, but roughly unaffected by loss of *Nrf1α^−/−^* or *Nrf2^−/−^*, no matter whether its basal expression of p-AKT was up-regulated in *Nrf1α^−/−^* cells or down-regulated in *Nrf2^−/−^* cells (Figure 4H, *h2*). Also, no significant changes in the PI3K and PTEN were observed in CDDP-intervened *Nrf1α^−/−^*cells, but the latter PTEN appeared to be more sensitive to CDDP in *Nrf2^−/−^* cells (Figure 4H, *h3*). In addition, p65 (involved in the NFκB signaling response) was basally enhanced in *Nrf1α^−/−^*or *Nrf2^−/−^* cells, but also almost unaffected by CDDP (Figure 4H, *h5*).

Since *Nrf1α^−/−^* cells gave rise to considerably higher mRNA expression of *CDH1*, *CDH2* and *MMP9* (Figure 4F), further Western blotting analysis was conducted to verify whether the loss of *Nrf1α^−/−^* is much likely to enable for transactivation of those EMT-related genes. As anticipated, the results unveiled that the epithelial-prone protein CDH1 was almost completely abolished in *Nrf1α^−/−^* cells and also largely unaffected by CDDP (Figure 4H, *h6*), but it was sharply suppressed by this drug intervention of *WT* cells. By contrast, the mesenchymal-prone proteins CDH2 and ITGB4 were basally enhanced by loss of *Nrf1α^−/−^* and also inducibly up-regulated by CDDP (Figure 4H, *h7, h9)*, but the former CDH2 expression was abolished in *Nrf2^−/−^* cells and also unaffected by this drug. Furtherly, SNAIL1 was highly expressed in *Nrf2^−/−^* cells when compared with that of *WT* or *Nrf1α^−/−^* cell lines (Figure 4H, *h8*), but its inducible expression levels were modestly up-regulated by CDDP in all three cell lines. Taken altogether, it is inferable that Nrf1 and Nrf2 are differentially involved in different cellular signaling responses to CDDP.

### 3.6 Reprogramming of cell metabolomes by Nrf1α^−/−^ or Nrf2^−/−^ in their metabolic responses to CDDP

Since discovery of the Werburg effect, it is generally accepted that the reprogramming of cell metabolism plays a crucial role for meeting the excessive demands for energy during cancer progression [36, 37]. This process also includes some key modifications within a variety of metabolic pathways (e.g., glycolysis, TCA, glutamine cleavage). Herein, the untargeted metabolomics analysis of different genotypic cell lines was performed in order to identify whether and/or which certain precise metabolic pathways are reprogrammed by the loss of *Nrf1α^−/−^* or *Nrf2^−/−^* in response to CDDP. As anticipated, 4867 of differential metabolites were screened by log_2_[fold changes] ≥ 1.5, based on the average expression levels of examined metabolites in *WT* cells. Amongst them, 896 metabolites were further selected with two opposite expression trends in *Nrf1α^−/−^*and *Nrf2^−/−^* cell lines (probably leading to different cancer cell fates [6],[5]).

As shown in Figure 5A, such 896 metabolites were enriched with the relevant KEGG pathways by using the MetaboAnalyst4.1 database. Notably, both metabolic pathways of *D*-glutamine and *D*-glutamate were displayed at the first place, along with *L*-glutamine as an essential compound prompted (Figure 5B). By the KEGG mapping of those metabolic pathways, *L*-glutamine expression was evaluated to be the highest metabolite in *Nrf1α^−/−^* cells to 15-fold changes when compared with that of being measured from *WT* or *Nrf2^−/−^* cell lines (Figure 5C). One of the *L*-glutamine metabolites, 2-oxo-glutarate (2-OG, as a key complement to the TCA cycle, which is linked to the electron transport chain and thus closely associated with ROS production [38, 39]), was also 2-5-fold higher than equivalents in *WT* and *Nrf2^−/−^* cells, whilst the another metabolite, 5-oxo-*D*-proline, was considerably lower in *Nrf2^−/−^* when compared to the counterpart of *WT* or *Nrf1α^−/−^* cells. Accordingly, the *D*-glutamate expression was less abundant, especially in *Nrf2^−/−^* cells (Figure 5C). This, taken together with a prior report that intracellular ROS influence the 2-OG-to-succinate ratio and the TCA cycle by regulating glutamine metabolism [40], suggests that the catalysis of putative substrate enzymes (and encoding genes) is critical for the biochemistry fate of 2-OG, which is also determined in the contexts within certainly unbalanced or homeostatic environments.

**Figure 5.**
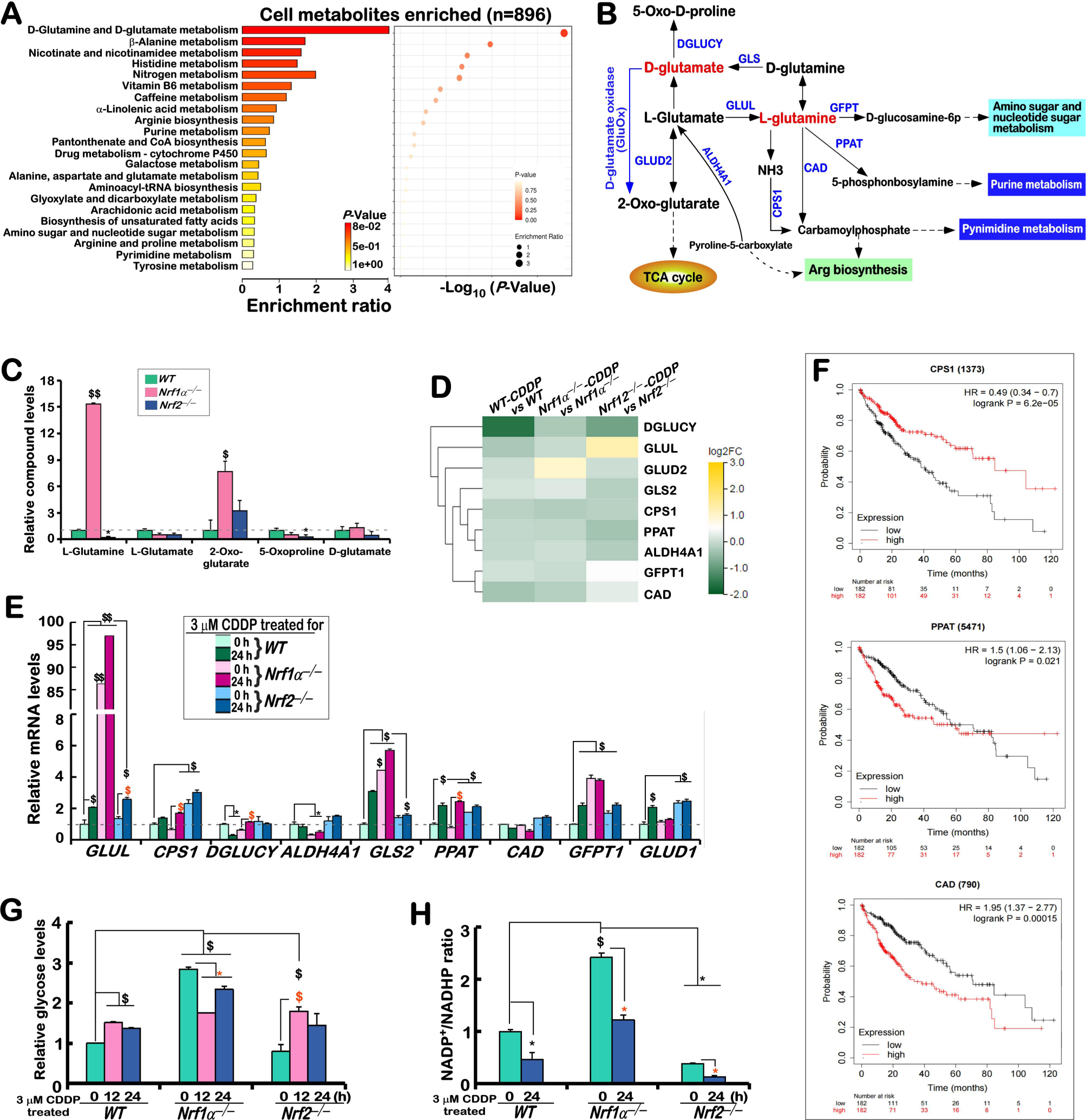
Reprogramming of cell metabolomes by Nrf1α^−/−^ or Nrf2^−/−^ in their metabolic responses to CDDP. **(A)** Twenty-two metabolic pathways enriched by 896 of differential metabolites with two opposite expression trends in *Nrf1α^−/−^* and *Nrf2^−/−^* cell lines, which were selected by using the online MetaboAnalyst4.1 database, among which the difference in the *D*-glutamine metabolism pathway is evaluated to be the most significant (*p* < 0.001). **(B)** A schematic of the *D*-glutamine metabolic pathway (adopted from the KEGG database), with those representative metabolites (in red) of significant differences in each of cell lines. The relevant enzymes are denoted (in blue), with both direct and indirect promoted relationships (which were represented by solid or dashed lines, respectively). **(C)** The contents of those key metabolites arising from the *D*-glutamine metabolic pathway were shown graphically as fold changes (mean ± SD, n = 3), with significant increases ($, *p* < 0.05; $$, *p* < 0.01) and significant decreases (\**p* < 0.05), which were statistically analyzed when compared with *WT* controls. **(D)** A heat-map shows differential expression profiling of those genes encoding key enzymes within the *D*-glutamine metabolism pathway of examined cell lines (*WT*, *Nrf1α^−/−^*and *Nrf2^−/−^*), that had been intervened for 24 h with CDDP or not, before being subjected to the transcriptome sequencing. **(E)** The basal and CDDP-stimulated expression levels of the above-described enzyme-encoding genes were validated by real-time qPCR in *WT*, *Nrf1α^−/−^* and *Nrf2^−/−^* cell lines, that had been intervened for 24 h with CDDP or not. The results were shown as fold changes (mean ± SD, n = 3×3), which are representative of at least three independent experiments being each performed in triplicates. Significant increases ($, *p* < 0.05; $$, *p* < 0.01) and significant decreases (\**p* < 0.05) were statistically analyzed when compared to *WT* controls. Some symbols “$ or *” indicate significant differences in *Nrf1α^−/−^*or *Nrf2^−/−^* cells compared to their respective *t0* after CDDP intervention. **(F)** Three key enzymes CPS1, PPAT and CAD of the *D*-glutamine metabolic pathway were manifested with a significant correlation to the survival period of patients with liver cancer (obtained from the online Kalpan-Meier plotter database). **(G,H)** The content of intracellular glucose (***G***) and the ratio of NADP^+^/NADPH **(*H*)** were determined in *WT*, *Nrf1α^−/−^* and *Nrf2^−/−^*cell lines, that had been intervened with CDDP for 0, 12 or 24 h. Each group of samples was repeated in at least three composite wells. The measured absorbance (*OD*) values were then statistically analyzed and graphically shown as mean ± SD (n = 3×3), along with significant increases ($, *p* < 0.05) and significant decreases (\**p* < 0.05; \*\**p* < 0.01), were statistically analyzed when compared with *WT*t0, respectively. Additional symbols “$ or *” indicate significant differences in *Nrf1α^−/−^*or *Nrf2^−/−^* cell lines compared to their respective *t0* value measured after CDDP intervention.

To clarify this, we further investigate changes in the basal and CDDP-inducible expression of those key genes involved in the above-mentioned metabolic pathway, by scrutinizing the transcriptome data (Figure 5D), followed by validation by real-time qPCR (Figure 5E). The results unraveled that basal expression levels of *GLUL*, *GLS2*, and *GFPT1* were markedly up-regulated in *Nrf1α^−/−^* cells, while *CPS1*, *DGLUCY*, *ALDH4A1* and *PPAT* were basally down-regulated by such loss of *Nrf1α^−/−^*, when compared to their counterparts of *WT* cells (Figure 5E). The intervention of *Nrf1α^−/−^* cells with CDDP caused substantial inducible increases in the mRNA expression levels of *GLUL*, *CPS1*, *GLS2, DGLUCY* and *PPAT* to certainly higher extents, but *ALDH4A1, CAD, GFPT1* and *GLUD1* were almost unaltered. In *Nrf2^−/−^* cells, basal expression levels of *ALDH4A1, CPS1, PPAT* and *GLUD1* were, conversely, up-regulated, but all other examined genes except *GLUL* were almost unaffected by CDDP (Figure 5E). By contrast, the intervention of *WT* cells with CDDP enabled evident induction of *GLS2, PPAT, GFPT1* and *GLUD1*, besides *GLUL*, but only *DGLUCY* was down-regulated by this drug. Next, by the Kalpan-Meier’s plotting of such key genes on the survival curves of patients with liver cancer, it was unveiled that the high expression of only *CPS1* and *PPAT* amongst the above-mentioned genes enables the patient survival to be significantly prolonged (Figure 5F), but the low expression of *CAD* can also prolong the survival of those patients (*lower panel*). In addition, no significant differences between all other examined gene expression status and the patients’ survival should also be noted herein.

From the above evidence showing that the expression levels of *GLUL* was up-regulated to different extents in examined three cell lines, particularly after CDDP intervention, it is hence inferable that it should be a key gene required for this metabolism to accelerate the conversion of *L*-glutamate to *L*-glutamine, leading to a yield of the massive metabolite 2-OG. Such being the case, 2-OG is held as a main resource to continuously replenish the TCA cycle and accelerate gluconeogenesis, in order to provide the energy and nutrients required for the proliferating cells. As expected, the relative glycose content (Figure 5G) and metabolism-coupled NADP^+^/NADPH ratio (Figure 5H) were substantially augmented in the proliferating *Nrf1α^−/−^* cells, but strikingly reduced in slowing *Nrf2^−/−^* cells. Notably, the intervention of *Nrf1α^−/−^* cells with CDDP caused an obvious decrease in the glycose content (possibly arising from gluconeogenesis), which remained to be modestly higher than those levels obtained from this drug stimulation of *WT* or *Nrf2^−/−^* cell lines (Figure 5G). However, the ratio of NADP^+^/NADPH was repressed to certainly lower extents by CDDP in all three examined cell lines (Figure 5H). This implies that the redox metabolism related to NADP^+^/NADPH is likely reprogrammed by CDDP intervention, and its nuances may be varying with the loss of *Nrf1α^−/−^* or *Nrf2^−/−^*.

To clarify which the precise metabolic pathways are reprogrammed by loss of *Nrf1α^−/−^* or *Nrf2^−/−^* in response to CDDP, the relevant transcriptome sequencing data were further parsed herein. As shown in Figure S2A, much more genes involved in the glycose, lipid, amino acid metabolisms were found in CDDP-intervened *Nrf2^−/−^*cells, but relatively less alterations of such genes were detected in CDDP-treated *Nrf1α^−/−^* cells, when compared to *WT* cells. Of note, CDDP intervention of *WT* cells caused up-regulation of those key genes converged in the fatty acid synthesis and the TCA cycle, but other genes responsible for the amino acid transport were significantly down-regulated. In *Nrf2^−/−^* cells, CDDP-stimulated differential expression genes (DEGs) were discretely spread in general matter metabolism, of which significantly up-regulated or down-regulated genes were primarily required for the gluconeogenesis, fatty acid synthesis and oxidation, and amino acid synthesis and transport, whereas relatively less changed genes are mainly involved in the glycolysis and TCA cycle (Figure S2A). Therefore, it is inferable that such metabolism programming of *Nrf2^−/−^* cells is likely attributable to induction of the remnant Nrf1 by CDDP. This notion was also supportively evidenced by further observations, revealing that relatively less intervened effects of CDDP on those metabolism genes, which were focally limited to some lipid metabolism, in its treated *Nrf1α^−/−^* cells (Figure S2A), although Nrf2 was aberrantly accumulated by the loss of *Nrf1α^−/−^* (Figure 1, D to I).

Subsequently, CDDP-stimulated mRNA expression of key genes involved in the glycose and lipid metabolisms were further corroborated by real-time qPCR (Figure S2, B & C). The results uncovered that CDDP intervention of all three examined cell lines enabled different extents of up-regulation of *GLUT1* and *GLUT4* (both essential for glucose transport), albeit their basal levels had been enhanced by such loss of *Nrf1α^−/−^* or *Nrf2^−/−^* (Figure S2B). Both *HK1* and *HK2* (encoding the rate-limiting enzymes for glycolysis) were up-regulated by CDDP in *Nrf1α^−/−^* cells, but down-regulated by this drug in *WT* cells, aside from that was roughly unaffected in *Nrf2^−/−^* cells. Intriguingly, it was found that all those critical genes for the lipid metabolism were not induced by CDDP, but instead mostly reduced by this drug intervention of examined cell lines, along with an exception of a few genes being unaffected, no matter whether their basal expression levels were enhanced or repressed in untreated *Nrf1α^−/−^* or *Nrf2^−/−^* cell lines (Figure S2C). At the protein levels, CDDP intervention of *WT* cells only stimulated modest increases in the expression of GLUT1, HK2 and LDHA (for glycose metabolism, Figure S2D), but apparently induced the expression abundances of ACCa and SCD (both essential for lipid metabolism, Figure S2E). In *Nrf1α^−/−^* cells, the abundances of GLUT1, HK1, HK2, and LDHA were certainly induced by CDDP (Figure S2D). Also, basal and CDDP-stimulated expression levels of SREBP2 were substantially elevated by this intervention of *Nrf1α^−/−^* cells, all other examined lipid metabolism-related proteins were down-regulated to different extents (Figure S2E). Similarly, the expression of such molecules required for lipid metabolism (including SREBP2) was all mostly diminished by CDDP in *Nrf2^−/−^* cells (Figure S2E), but conversely most of the examined glycose metabolism proteins were modestly up-regulated by this drug (Figure S2D). Taken altogether, these results demonstrate that both Nrf1 and Nrf2 are differentially contributable to the metabolism reprogramming response to CDDP.

### 3.7 In vivo malgrowth of xenograft tumors was effectively intervened by CDDP via Nrf1-dependent and -independent pathways

Our previous studies had shown that *in vivo* malgrowth of *Nrf1α^−/−^*-derived xenograft tumors was significantly incremented in nude mice, whereas *Nrf2^−/−^*-derived tumor growth was substantially blocked [6, 30]). Herein, we continuously investigated the *in vivo* intervening effect of CDDP on the malignant progression of *Nrf1α^−/−^*-derived, rather than *Nrf2^−/−^*-derived tumors in mouse xenograft model. As illustrated in Figure 6 (A & B), it was confirmed that *Nrf1α^−/−^* transplanted tumors were rapidly grown, with approximately 4.3 times larger in size than that of *WT* tumor. CDDP intervention was administrated on the 4^th^ day of subcutaneous tumor formation and then its inhibitory effect of this intervention on xenograft tumors was seemingly observed to be roughly unaffected by the presence of *Nrf1α* (in *WT*+CDDP group) or its absence (in *Nrf1α^−/−^*+CDDP group). However, histopathological examination revealed obvious coagulative necrosis of CDDP-intervened tumors, which were more severely in the *WT*+CDDP group than *Nrf1α^−/−^*+CDDP group (Figure 6C), even although their tumor volumes and weights were all strikingly repressed by this drug intervention (Figure 6, D & E). Further Tunnel staining unraveled that malgrowth of these tumors was inhibited by CDDP through apoptosis (Figure 6F), which is thereby contributable to its active functioning as a mature anticancer agent [41]. However, there appeared to exist subtle nuances in tumor volume and weight after CDDP intervention, which were manifested with a stronger inhibitory effect of this drug on *WT* tumors than its effect on the *Nrf1α^−/−^*-deficient tumor (*cf.* 0.3 for *WT*+CDDP *vs WT* with 0.38 for *Nrf1α^−/−^*+CDDP *vs Nrf1α^−/−^*). This indicates that CDDP was, perhaps, more effective on the *WT* tumors in the presence of *Nrf1α* (Fig. 6D, E). This is supported by another closer observation, showing that a partial recovery of such *Nrf1α^−/−^*-deficient xenograft tumors from inhibition of CDDP occurred from the 8^th^ day after this drug was injected intraperitoneally (Figure 6A).

**Figure 6.**
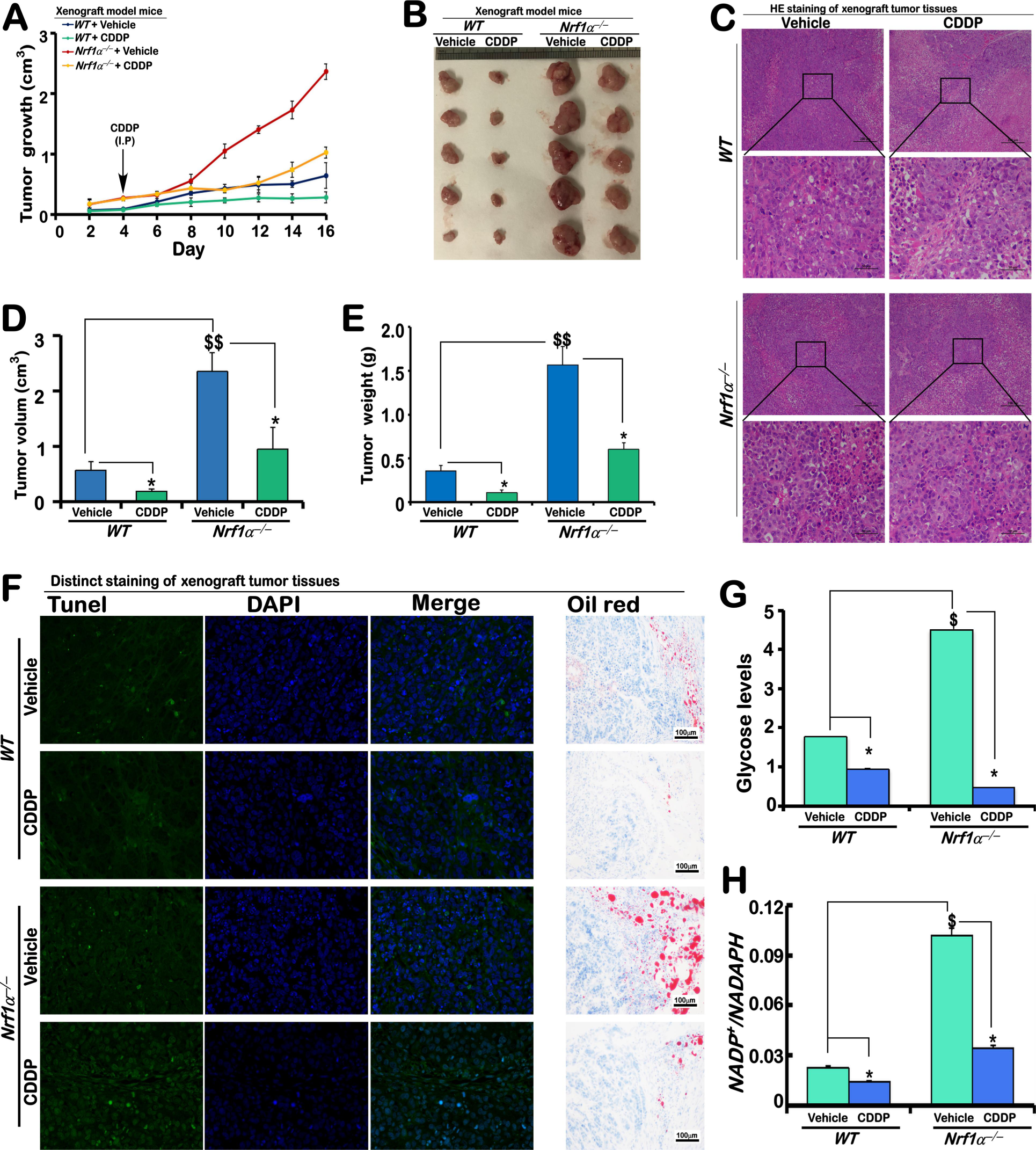
CDDP inhibits the *in vivo* malgrowth of xenograft tumors arising from *WT* and *Nrf1α^−/−^* cells in nude mice. **(A)** The *in vivo* malgrowth curve of xenograft tumors in distinct groups of nude mice, that were subcutaneously injected with human *WT* or *Nrf1α^−/−^* hepatoma cell lines, followed by intervention with CDDP (10 mg/k) or vehicle (i.e., 0.9% physiological saline). **(B)** The xenograft tumors were removed after all the model mice were euthanized on the 16^nd^ day after CDDP intervention and then exhibited herein. **(C)** The model tumor tissues were subjected to the histopathological examination by routine HE staining, and then relevant images were obtained from microscopy. **(D,E)** The volumes (***D***) and weight **(*E*)** of the above xenograft tumors were determined as shown graphically. **(F)** The staining of TUNEL and oil red O was subjected to evaluating cell apoptosis and lipid storage in xenograft tumor tissues. **(G,H)** The content of glycose and the ratio of NADP^+^/NADPH within tumor tissues were detected, which were performed at least in triplicates. The obtained absorbance values are statistically determined and shown graphically, with significant increases ($, *p* < 0.05) and significant decreases (\**p* < 0.05), relative to the control value of each group as indicated.

In the meantime, the oil red O staining demonstrated that CDDP caused an apparent reduction in the lipid deposition by fatty acid metabolism and/or lipid droplet formation within indicated tumor tissues, particularly in CDDP-intervened *Nrf1α^−/−^*group (Figure 6F). The glycose content and the NADP^+^/NADPH ratio were alleviated simultaneously by CDDP in those intervened tumor tissues, which also appeared to be more effective (by 2 to 4-fold reduction) in the *Nrf1α^−/−^* +CDDP group (Figure 6, G &H). This indicates a higher degree of cancer malignancy is augmented if *Nrf1α* is lost, when compared to its presence in *WT* tumors, but when CDDP induces apoptosis of tumor cells, the consequent requirement for energy is further reduced to a certain lower extent.

### 3.8 The intervened effect of CDDP on the tumor tryptophan metabolism pathways is monitored by Nrf1

Herein, we performed an untargeted metabolome analysis of xenograft tumor tissues arising from *WT* and *Nrf1α^−/−^*-deficient cell lines. As shown in Figure 7A, the results revealed 2310 differential metabolites of between two types of tumor tissues, of which 303 metabolites were selected by calculating their different parameter of log_2_FC (fold changes) ≥ 2. Amongst them, 195 metabolites were up-regulated, whereas 108 metabolites were down-regulated in *Nrf1α^−/−^*-deficient tumor tissues (as compared to the *WT* control) (Figure 7A, *a2*). By using the online MetaboAnalyst4.1 database, such significantly differential metabolic compounds of 303 were enriched based on the KEGG pathway, revealing that amongst the metabolic pathway of xenograft tumor tissues, the first place is taken predominantly through the tryptophan metabolism (Figure 7B).

**Figure 7.**
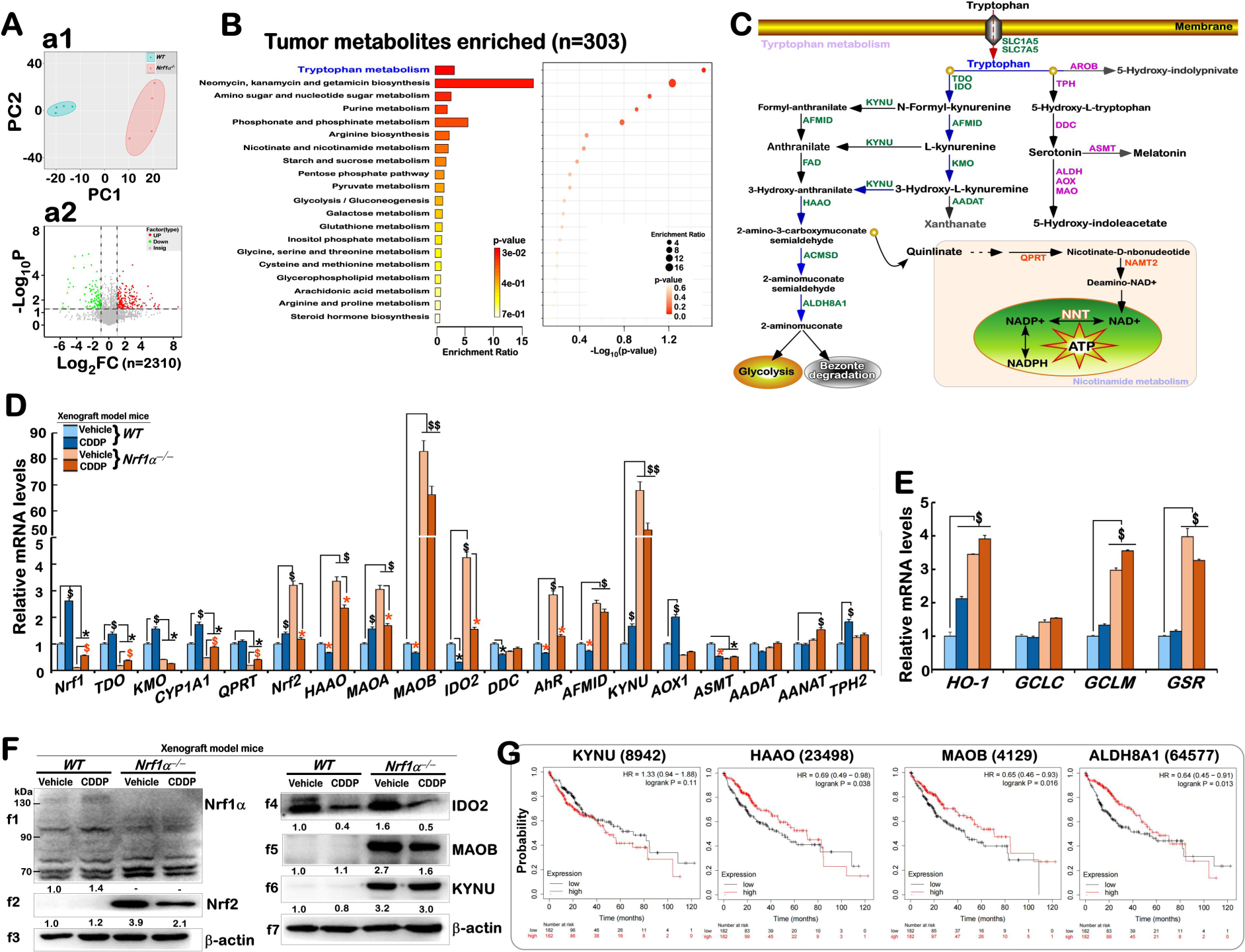
The intervened effect of CDDP on tryptophan metabolism pathways of xenograft tumors. **(A)** Untargeted metabolome analysis of the whole-tumor metabolites in *WT* or *Nrf1α^−/−^*-deficient tissues (which was performed at least four duplicates in each group). Of note, principal component analysis (PC) was shown (***a1***), and differential expression compounds **(*a2*)** was illustrated as the volcano plot (Red, up-regulated; Green, down-regulated; Gray, no difference) between the two tissue metabolic species. **(B)** Differential metabolites between *WT* and *Nrf1α^−/−^*-deficient tumor tissues were mainly enriched in the tryptophan metabolism pathway being taken at the first place. **(C)** A schematic diagram shows tryptophan transport and its metabolic pathway (adapted from the online KEGG pathway database). The kynurenine (kyn) metabolic branch in tryptophan metabolism was arrowed in blue. The enzymes in the kyn branch (green), the enzymes in the serotonin biosynthesis (pink), and the other enzymes in the nicotinate acid metabolic pathway (orange) mediated by quinolate at the end of the kyn metabolism were indicated herein. **(D,E)** Real-time qPCR analysis was subjected to detecting the effects of CDDP intervention on transcriptional expression of those key enzymes (e.g., *Nrf1, TDO, KMO, CYP1A1, QPRT, Nrf2, HAAO, DDC, AhR, AFMID, KYN, AOX1, ASMT, AADAT, AANAT, TPH2*) in tryptophan metabolism pathway (*D*) and the expression levels of antioxidant genes (involved in the GSH-based redox signaling) (*E*) in different tumor tissues after intervention by CDDP or vehicle. The resulting data were shown as fold changes (mean ± SD, n = 3 × 3), which are representative of at least three independent experiments being each performed in triplicates. Significant increases ($, *p* < 0.05; $$, *p* < 0.01) and significant decreases (\**p* < 0.05), were statistically analyzed when compared to *WTvehicle*, respectively. Additional symbols “$ or *” indicate significant differences in *Nrf1α^−/−^* tumors as compared to their *vehicle* controls. **(F)** Western blotting analysis was subjected to detecting the protein expression of Nrf1, Nrf2, IDO2, MAOB and KYNU in different tumor tissues. The intensity of those immunoblots was also quantified by the Quantity One 4.5.2 software, and showed below the indicated protein bands. **(G)** Those relationships of *KYNU, HAAO, MAOB*, and *ALDH8A1* in the tryptophan metabolism pathway to the survival rate of patients with liver cancer were obtained from the online Kalpan-Meier plotter database.

As shown schematically in a proposed model (Figure 7C), tryptophan is one of the amino acids essential for protein synthesis and also metabolically generates more biologically active catabolic metabolites, which include serotonin and other metabolites in the kynurenine (Kyn) pathway (*via* those key enzymes encoded by *IDO, KYNU, HAAO, QPRT* [42]. Clearly, the normal tryptophan metabolites are directly regulated by tissue-specific expression of tryptophan metabolizing enzymes [43], but altered expression of these enzymes in cancer leads to an increase in specific metabolic byproducts contributing to tumor development [44] [45]. Thus, we here examined whether such key enzymatic genes were influenced by CDDP in its intervened tumor tissues. As expected, the results unraveled significant differences in the mRNA expression of major genes in this pathway between two groups of tumor tissues from CDDP- and vehicle-intervened xenograft mice. In CDDP-intervened *WT* tissues, the expression of *TDO, KMO, CYP1A1, MAOA, KYNU, AOX1* and *TPH2* were up-regulated to varying degrees by this drug, in which *Nrf1* was also significantly up-regulated, whereas *Nrf2* was only modestly induced (Figure 7D). By contrast, *MAAO, MAOB, IDO2, DDC, AHR, AFMID* and *ASMT* were down-regulated by CDDP in *WT* tumor tissues. Just the opposite of most genes happened in *Nrf1α^−/−^*-deficient tumors. Basal mRNA expression levels of *TDO, KMO, CYP1A1, QPRT, DDC, AOX1,* and *ASMT* were markedly blocked by the loss of *Nrf1α^−/−^* (Figure 7D), and even upon stimulation by CDDP, they were almost unaffected or only marginally induced close to the basal levels of *WT-vehicle* tumors. Yet, it is noteworthy that *Nrf2, HAAO, MAOA, MAOB, IDO2, AHR, AFMID,* and *KYNU* were substantially augmented at their basal expression, but rather diminished or even abolished by CDDP in *Nrf1α^−/−^*-deficient tumors (Figure 7D). With very few exception, only *AANT* was slightly induced by this drug, while *AADAT* and *TPH2* were unaltered. In addition, *HO-1, GCLC, GCLM* and *GSR* were significantly expressed in *Nrf1α^−/−^*-deficient tumors, as consistent with the aforementioned cell-level experiments, but they appeared to be marginally changed or largely unaffected by CDDP (Figure 7E). However, only *HO-1* was significantly induced in CDDP-intervened *WT* tumor tissues.

Further immunoblotting analysis revealed that the protein expression levels of Nrf1, Nrf2, IDO2 and MAOB appeared to be roughly consistent with their above-described mRNA expression changes (Figure 7F). Albeit basal abundances of Nrf2, IDO2, MAOB, and KYNU were substantially elevated in *Nrf1α^−/−^*-deficient tumor tissues as compared to *WT-vehicle* tumors, they were quite down-regulated by CDDP intervention, along with an exception of KYNU that appeared to be less changed by this drug (Figure 7F). Moreover, the relationships of *KYNU, HAAO, MAOB* and *ALDH8A1* involved in the tryptophan metabolism pathway to the survival of patients with liver cancer were obtained from the Kalpan-Meier plotter database (Figure 7G). Of note, the survival rate of such hepatoma patients with *KYNU* was different from similar patients with *HAAO, MAOB* and *ALDH8A1*. Overall, these collective results indicate that Nrf1 is likely to monitor the basal expression levels of Nrf2 and those key genes in the tumor tryptophan metabolism pathways, so that both CNC-bZIP factors can make certainly differential or even opposing contributions to the intervened effect of CDDP on liver cancer.

### 3.9 Differential regulation of target genes by Nrf1 and Nrf2 in distinct genotypic cellular responses to CDDP

In order to clearly delineate CDDP-triggered signaling towards target genes regulated by Nrf1 or Nrf2 alone or together, we scrutinized the transcriptome sequencing data by in-depth comparative analysis of those indicated cell lines that had been intervened with CDDP. As shown in Figure 8A, the number of CDDP-upregulated genes were more than that of its down-regulated genes in three different cell lines, especially upon loss of *Nrf2*, which led to up-regulation of 65 genes that were eight times of the number of its downregulated genes. This was fully consistent with the volcano results from *Nrf2^−/−^*(Figure 8B). Intriguingly, by intersecting different genotypic cell lines intervened by CDDP, it was found that none of the putative joint genes functioning in all examined cell lines existed (through bioinformatic analysis of |Log_2_FC| ≥ 1.0) (Figure 8C). This implies that CDDP alone cannot directly regulate possibly inducible genes, except that it is only enabled for significant stimulation of those target genes regulated by Nrf1 and/or Nrf2 (Figure 8C).

**Figure 8.**
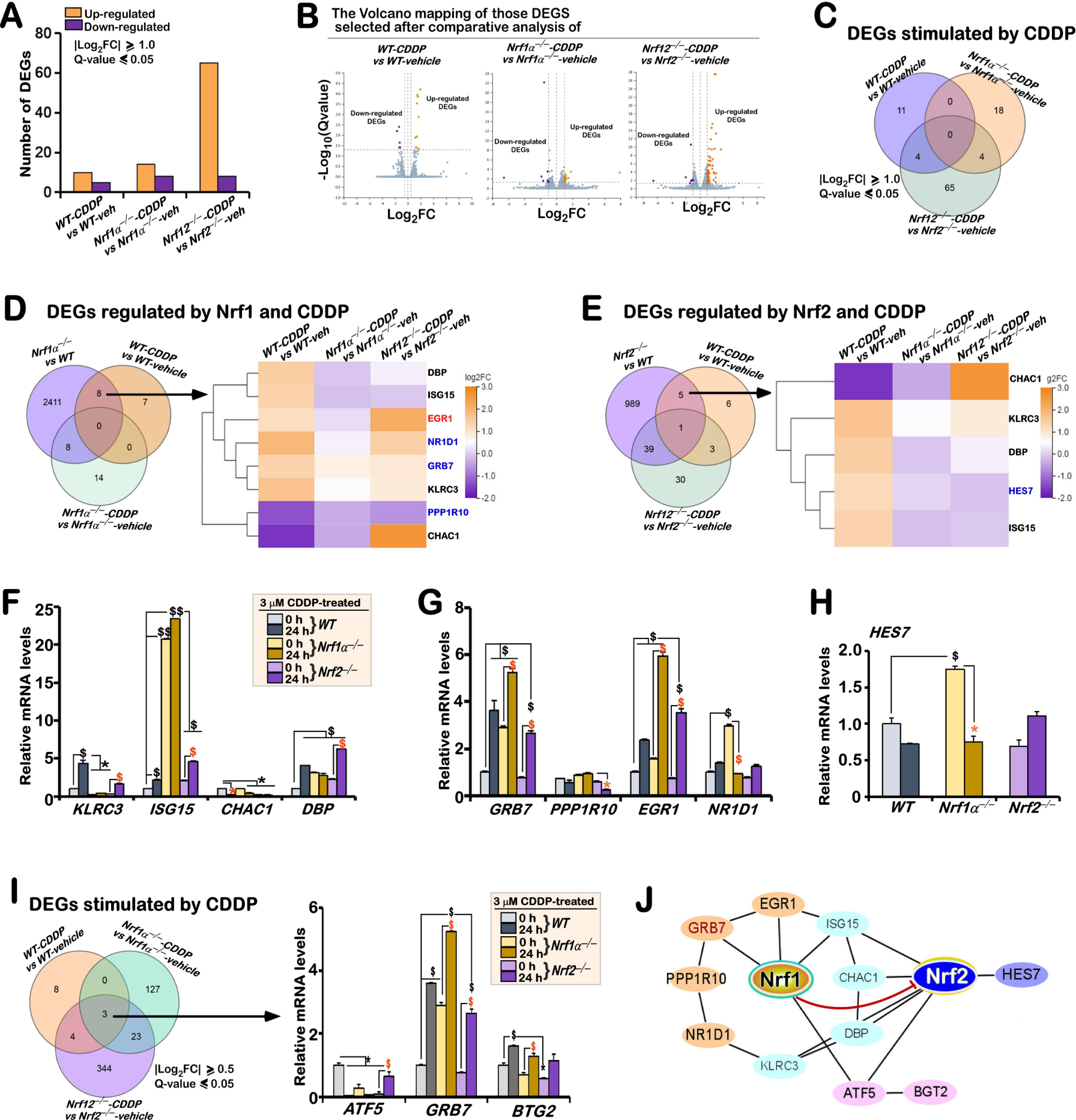
The pharmacological effect of CDDP on hepatoma cells is mainly achieved by stimulating target genes regulated by *Nrf1*. **(A)** Distinct regulation of those differential expression genes by CDDP in *WT*, *Nrf1α^−/−^* and *Nrf2^−/−^* cell lines was further analyzed. The intervened effects of CDDP on such genes were statistically determined relative to their corresponding vehicle controls, and then shown graphically. **(B)** Those differentially regulated genes were illustrated by distinct volcano plots. Orange represents gene up-regulation by CDDP, purple denotes gene down-regulation by CDDP, whereas gray shows no-significant differences. **(C)** The Venn plot shows distinct cell group-intersected genes stimulated by CDDP, which were selected mainly by calculating the parameter of CDDP-induced fold changes (i.e., log2FC ≥ 1.0). **(D,E)** The Venn plot shows those genes mediated mainly by Nrf1 (*D*) or Nrf2 (*E*) after CDDP intervention, and their differential expression changes were also further presented within distinct heat-maps, after being comparatively analyzed in the three pairs of differently treated cell lines. **(F-H)** Real-time qPCR analysis of differential gene expression stimulated by CDDP (*F*), and also regulated by *Nrf1* **(*G*)** or *Nrf2* **(*H*)**. The resulting data were shown as fold changes (mean ± SD, n = 3 × 3), which are representative of at least three independent experiments being each performed in triplicates. Among them, significant increases ($, *p* < 0.05; $$, *p* < 0.01) and significant decreases (\**p* < 0.05), were statistically determined, when compared with their corresponding *WT* controls. Additional symbols “$ or *” indicate significant differences in *Nrf1α^−/−^*or *Nrf2^−/−^* cell lines, relative to their *T0* values after CDDP intervention. (I) Those genes were selected by calculating the parameter log2FC ≥ 0.5 of CDDP-induced fold changes, and shown in the Venn plot. The differential expression levels of CDDP-specifically induced genes were determined by real-time qPCR in *WT*, *Nrf1α^−/−^* and *Nrf2^−/−^*cell lines that had been intervened with CDDP for 0 or 24 h. The resulting data were statistically analyzed as described above. (J) A schematic representation of the signaling gene-regulatory network mediated by Nrf1 and/or Nrf2 in response to CDDP. Of note, Nrf2 is tightly governed by Nrf1, primarily occurring at their protein levels as described elsewhere [6, 24, 30, 54].

Next, further bioinformatic comparative analysis unraveled CDDP-stimulated expression of 8 genes mediated by NRF1α, and different expression levels of such genes including *KLRC3*, *GRB7*, *PPP1R10*, *ISG15*, *CHAC1*, *DBP*, *EGR1* and *NR1D1* were shown in the heatmap (Figure 8D), in addition to other 8 genes independent of Nrf1α. By contrast, only 5 of CDDP-stimulated genes *KLRC3*, *ISG15*, *CHAC1*, *DBP*, and *HES7* was regulated by Nrf2, but 39 of other genes stimulated by CDDP were independent of Nrf2 (Figure 8E). The transcriptome sequencing uncovered significant changes in CDDP-stimulated expression of most genes (possibly regulated by Nrf1 and/or Nrf2) in *WT* and *Nrf2^−/−^* cells, but they were unchanged or even down-regulated by this drug in *Nrf1α^−/−^* cells (Figure 8, D & E). Among them, only 4 genes *KLRC3*, *ISG15*, *CHAC1* and *DBP* with differential expression responses to CDDP were inferably co-regulated by both Nrf1 and Nrf2 (Figure 8F). Subsequently, real-time PCR revealed that intervention of *WT* cells by CDDP led to significant increases in the expression of *KLRC3* (a lectin-like transmembrane receptor signaling in tumor immune response [46]), *ISG15* (a ubiquitin-like protein that mediates post-translational modification and protein processing in various biological processes [47]) and *DBP* (a D-box-binding PAR bZIP transcription factor to *trans*-activate *Albumin* and *CYP2A4/5* [48]). Of note, basal expression of *KLRC3* was almost completely abolished by loss of *Nrf1α^−/−^* or *Nrf2^−/−^* when comparison to *WT* controls (Figure 8F), but it was still partially induced by CDDP in *Nrf2^−/−^* cells, while it was rather unaffected by this drug in *Nrf1α^−/−^* cells. By contrast, basal expression of both *ISG15* and *DBP* was markedly enhanced by the loss of *Nrf1α^−/−^* or *Nrf2^−/−^*, but the former *ISG15* was further augmented by CDDP in either of deficient cells, whereas the latter *DBP* was up-regulated only by this intervention of *Nrf2^−/−^*, but not *Nrf1α^−/−^*, cell lines (Figure 8F). Conversely, *CHAC1* (glutathione-specific γ-glutamine transferase 1, that regulates the intracellular redox balance in UPR [49]) was diminished and even abolished by CDDP in three examined cell lines, particularly in *Nrf2^−/−^* cells. Collectively, these indicate that these four genes (putatively co-regulated by Nrf1 and/or Nrf2) appear to be more susceptible to induction by CDDP in both cell lines of *WT* and *Nrf2^−/−^* (with the remnant Nrf1 expressed), rather than *Nrf1α^−/−^* cells (albeit hyperactive Nrf2 accumulated). This also implies a vital role for Nrf1 in mediating these gene regulatory responses to CDDP.

Additional 4 genes such as *GRB7*, P*PP1R10*, *EGR1*, and *NR1D1* were postulated to be Nrf1-specific responsive to CDDP and then further validated by real-time qPCR (Figure 8G). The results revealed significant increases in basal and CDDP-stimulated expression of *GRB7* (growth factor receptor bound protein 7, enabling activation of the AKT signaling to promote the G1/S phase transformation [50]) and *EGR1* (early growth response 1, acting as a C2H2 zinc finger transcriptional regulator of target genes required for differentiation and mitogenesis [51]) in *Nrf1α^−/−^* cells. Both *GRB7* and *EGR1* were also up-regulated to certain higher extents by CDDP in *WT* and *Nrf2^−/−^* cell lines. However, almost no obvious changes in the expression of *PPP1R10* (protein phosphatase 1 regulatory subunit 10, directly interacting with the lipid-binding C2 domain of PTEN so as to be sequestered in the nucleus) were determined in all three examined cell lines (Figure 8G). Furtherly, *NR1D1* (a member of nuclear receptor subfamily 1D, acting as a transcriptional repressor in metabolic pathways in a heme-dependent manner[52]) was basally up-regulated in *Nrf1α^−/−^* cells, but rather seriously suppressed by CDDP to the untreated *WT* control levels, albeit it was only marginally altered or even unaffected in *WT* and *Nrf2^−/−^* cell lines (Figure 8G). In the meantime, among the putative Nrf2-specific responsive genes to CDDP, *HES7* (a bHLH transcription factor involved in the developmental Notch signaling pathway [53]) was found to be highly expressed in *Nrf1α^−/−^*cells and partially repressed in *Nrf2^−/−^* cells (when compared to the *WT* control) (Figure 8H), implying a relevance to the hyperactive Nrf2. Upon intervention by CDDP, the *HES7* expression was substantially diminished by the loss of *Nrf1α^−/−^* to a considerably low extent, similar to this drug-reduced expression level in *WT* cells, but the loss of *Nrf2^−/−^* resulted in a modest increased response to this intervention (Figure 8H).

Moreover, it was re-found by bioinformatic analysis of |Log_2_FC| ≥ 0.5 that three joint genes (i.e., *ATF5, GRB7, BTG2*) were intersected in all examined cell lines intervened by CDDP (Figure 8I, *left panel*). Aside from *GRB7* as described above, the expression of *BTG2* (an anti-proliferation regulator of the BTG/TOB family) was up-regulated by CDDP to varying extents within three examined cell lines (Figure 8I, *right panel*), albeit its basal levels were down-regulated by loss of *Nrf1α^−/−^* or *Nrf2^−/−^*. Also, basal expression of *ATF5* (as a key player in the mitochondrial unfolded protein response, i.e., UPR^mito^) was substantially diminished or completely abolished by loss of *Nrf1α^−/−^* and *Nrf2^−/−^*, respectively. By contrast, CDDP-stimulated expression of *ATF5* was also totally abolished in *WT* cells, in addition to *Nrf1α^−/−^*cells, but modestly increased relative to the basal level in *Nrf2^−/−^*cells. Taken together, these results demonstrate that Nrf1 and Nrf2 are differentially contributable to distinct effects of CDDP on those genes, because they appear to be more susceptible to CDDP intervention in *WT* cells (with the presence of both factors), even upon the absence of *Nrf2^−/−^*(with the remnant expression of Nrf1), rather than of *Nrf1α^−/−^*(albeit hyperactive Nrf2 accumulated).

### 3.10 Discrepant involvement of Nrf1 and Nrf2 in the drug resistance pathway to CDDP

To gain an insight into distinct contributions of Nrf1 and Nrf2 to mediating the drug resistant response to CDDP, we here examine their deficient effects on those key genes involved in the drug resistance pathway. The heatmap showed that specific loss of Nrf1α or Nrf2 caused distinct extents of decreases in most genes required for the putative drug resistance, except that a few of genes *GSTO2, POLH* and *XPA* were increased (Figure 9A). Rather, real-time qPCR revealed a significant distinction in the regulation of such key genes by Nrf1α from Nrf2 (Figure 9B). This is due to that fact that obvious upregulation of *ABCB1* (encoding a multiple drug-resistant glycoprotein P-gp to function as an ATP-binding cassette transporter), *REV3L, XPA* and *BRCA1* (the latter 3 genes required for DNA damage sensing and relevant repair) in *Nrf1α^−/−^* cells was accompanied by down-regulation of *POLH, MSH2, TOP2A, ATM, TP53* and *BCL2*. By contrast, strikingly opposing expression levels of these genes were determined in *Nrf2^−/−^* cells (Figure 9B), along with an exception that both *MDM2* and *MLH1* were up-regulated only in *Nrf2^−/−^* cells, while *GSTO2* was up-regulated only in *Nrf1α^−/−^* cells. In addition, high expression of *ATP7A* (ATPase copper transporting α) in the two deficient cell lines (Figure 9B), but another high-affinity copper transporter-encoding *SLC31A1* was markedly down-regulated in *Nrf2^−/−^* cells by comparison with its expression in *Nrf1α^−/−^* cells (Figure 9A). Further Western blotting analysis unraveled that high expression of P-gp was accompanied by hyperactive Nrf2 accumulation in *Nrf1α^−/−^* cells, but both were almost completely abolished in *Nrf2^−/−^* cells (Figure 9C, *c1 vs c8*). All other examined p53, Bcl2, H2AX and γ-H2AX were down-regulated in *Nrf1α^−/−^* cells, but also conversely up-regulated in *Nrf2^−/−^* cells, albeit their changed extents were varied within different cellular contexts (Figure 9C, *c2 to c5*). Collectively, these demonstrate distinct or even opposite contributions of Nrf1α and Nrf2 to monitoring putative drug resistance response, in which Nrf2 is endowed to play an irreplaceably predominant role, whereas Nrf1 is likely required for the DNA damage repair response. The latter notion is further supported by the ChIP-sequencing data, revealing a potent binding activity of Nrf1, rather than Nrf2, to the promotor regions of those relevant genes *H2AX, XPA, TOP2A,* and *MSH2* (Figure S3, adapted from https://www.encodeproject.org/).

**Figure 9.**
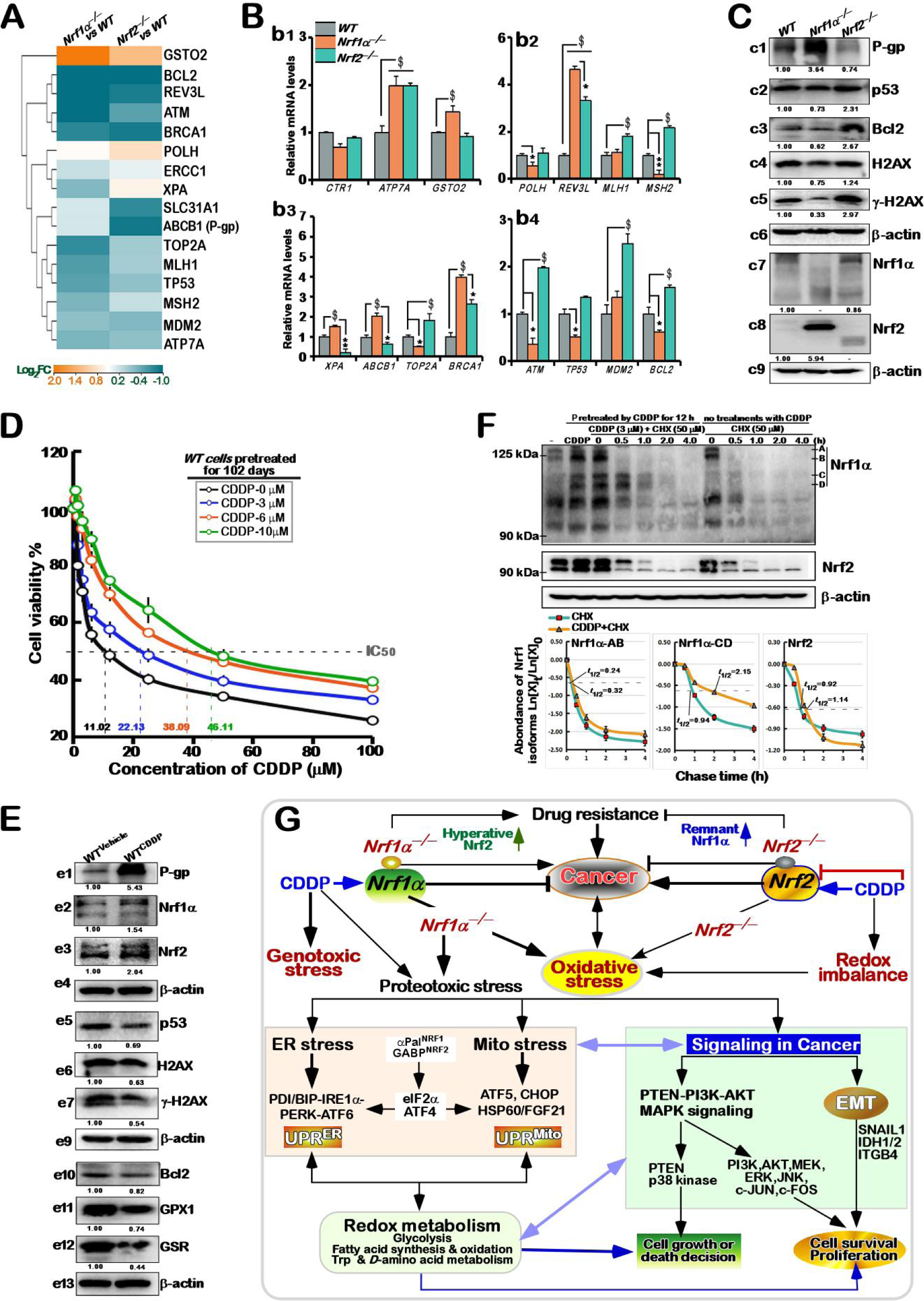
Distinct involvement of Nrf1 and Nrf2 in the drug resistance pathway to CDDP. **(A)** The heat-map shows changes of those differential expression genes in the putative drug resistance, which were determined in *Nrf1α^−/−^* or *Nrf2^−/−^* cell lines when compared to *WT* cells. **(B)** Basal mRNA expression levels of key genes involved in drug resistance pathway were validated by real-time qPCR analysis in *WT, Nrf1α^−/−^* or *Nrf2^−/−^* cell lines. The resulting data were shown as fold changes (*mean ± SD*, *n* = 3 × 3), which are representative of at least three independent experiments being each performed in triplicates. Significant increases ($, *p* < 0.05) and significant decreases (\**p* < 0.05; \*\**p* < 0.05) were statistically analyzed when compared with *WT* (measured at 0 h), respectively. **(C)** The basal protein expression abundances of key genes required for drug resistance were determined by Western blotting with relevant antibodies. The intensity of those immunoblots was also quantified by the Quantity One 4.5.2 software, and showed below the indicated protein bands. **(D)** Differential resistances of distinct cell populations to CDDP were evaluated by the viability of *WT* cells that had been pretreated with lower doses (0, 3, 6 or 10 μM) of this drug for 102 days before being intervened with distinct concentrations of CDDP (0 to 100 μM) for 24 h. The IC50 was calculated and also indicated herein. **(E)** The 10 μM CDDP-pretreated *WT* cells (i.e., *WT*^CDDP^) along with *WT*^Vehicle^ cells were subjected to Western blotting analysis of key genes involved in the drug resistance pathway. The intensity of those immunoblots was quantified as described above. **(F)** Distinct effects of CDDP on the stability of Nrf1 and Nrf2 were estimated in *WT* cells, which had been were treated with 50 µg/ml of cycloheximide (CHX) alone or plus 3 μM CDDP for indicated lengths of time in the pulse chase experiments, followed by visualization by Western blotting. The intensity of those interested immunoblots was quantified by the Quantity One 4.5.2 software. The resulting data are shown graphically, with distinct protein half-lives of longer or processed Nrf1, as well as Nrf2, after being calculated by a formula of Ln([A]t /[A]0, in which [A]t indicated a fold change (mean ± SD) in each of those examined protein expression levels at different times relative to the corresponding controls measured at 0 h (i.e., [A]0). **(G)** A proposed model is provided for a better understanding of distinct molecular mechanisms by which antioxidant transcription factors Nrf1 and Nrf2 are differentially or coordinately contributable to the anticancer efficacy of CDDP on hepatoma cells and even its drug resistance. Distinct cellular signaling responses to CDDP were induced in its intervention or chemoresistance.

In order to test the resistance of *WT* cells to CDDP, the incremental concentrations of this drug were added in the culture media, and then growth in different concentrations of this drug for 102 days (Figure 9D). The capability of such pretreated cell populations to resist against different doses of CDDP was evaluated by assaying their distinct viability. The results uncovered that the higher doses of CDDP (from 3 μM to 10 μM) were treated into those cell populations, the more potentiality of those cells to resist against subsequent intervention of this drug (from 3 μM to 100 μM) was developed (Figure 9D). The IC_50_ values of such distinct cell populations that had been pretreated for 102 days with distinct doses of CDDP (0, 3, 6, 10 μM) were determined to be 11.02, 22.13, 38.09 or 46.11 μM of this drug as intervened, respectively. Further Western blotting analysis of the above 10 μM CDDP-pretreated cell population (i.e., *WT^CDDP^*), along with *WT* cells pretreated with vehicle (i.e., *WT^V^*^ehicle^), revealed that highly-expressed abundance of P-gp in *WT^CDDP^* cells was accompanied by enhanced expression of Nrf2, as well by partially-increased expression of Nrf1 (Figure 9E, *e1 to e3*). Such being the case, all the other examined proteins p53, H2AX, γ-H2AX, Bcl2, GPX1 and GSR remained to be down-regulated to variously lower extents in *WT^CDDP^* cells, when compared to their counterparts in *WT^V^*^ehicle^ cells (Figure 9E, *e5 to e12*). Taken together, these results indicate distinct roles of Nrf2 and Nrf1 in the drug resistance and other cytoprotective defense against CDDP.

Lastly, distinct effects of CDDP on the stability of both Nrf1 and Nrf2 were also evaluated by cycloheximide (CHX)-pulse chase experiments (Figure 9F, *upper two panels*), followed by calculation of their protein half-lives as shown graphically (*lower panels*). The results unraveled no significant changes in the half-lives of the full-length Nrf1α-A/B (0.24 h *vs* 0.32 h) and Nrf2 (0.92 h *vs* 1.14 h) after being intervened with CDDP, but the half-lives of those processed Nrf1α-C/D proteins were strikingly prolonged by this drug from 0.94 h (=56 min) to 2.15 h (=129 min) (Figure 9F, *lower middle panel*). This finding demonstrates that CDDP significantly enhances stabilization of the active processed Nrf1α-C/D, rather than Nrf2, such that this mature Nrf1 factor can effectively regulate the transcriptional expression of those genes involved in CDDP-induced DNA damage response and relevant repair cascades, as well as other cytoprotective systems.

## 4. Discussion

Although CDDP has been widely administrated as an anticancer alkylating agent with dual functional groups to yield those redox-active substances with strong electrophilic properties and hence react with any nucleophilic molecules, it is less known about which (and how) electrophilic/antioxidant responsible transcription factors Nrf1 and Nrf2 modulate such redox signaling from cellular metabolism to gene reprogramming, and even adaptive reshaping after intervention with this drug. In this study, we have found: i) differential activation of Nrf1 and Nrf2 in mediating distinct cellular responses to CDDP; ii) substantial suppression of *in vivo* malgrowth of *WT*-rather than *Nrf1α^−/−^-*derived tumors by CDDP in the xenograft model mice, but a part of the lagged recovery from such repressing effect of this drug occurring only in *Nrf1α^−/−^* tumors (albeit hyperactive Nrf2 is retained); iii) significant resistance of *Nrf1α^−/−^* (more rather than *Nrf2^−/−^*) cell lines to the cytotoxicity (and genotoxicity) of this drug when compared to that of *WT* cells; and iv) distinct cellular and molecular mechanisms by which Nrf1 and Nrf2, as drug targets, are differentially or coordinately contributable to the anticancer efficacy of CDDP on hepatoma cells (and xenograft tumors), and also to this drug resistance (Figure 9G).

### 4.1 Differential activation of Nrf1 and Nrf2 in mediating distinct cellular responses to CDDP

Since as a redox-active electrophilic reagent, CDDP (and derivatives) can thus be enabled to covalently react any nucleophilic residues on a vast variety of biological molecules, particularly those macromolecules involved in cell components and life processes [55]. Such biochemical reactions should have resulted in directly destructive damages to primarily prototypic macromolecules (e.g., RNA, DNA, proteins) and other secondary harmful effects, in addition to redox imbalance and ensuing oxidative stress. Thereby, it is predictable that this electrophilic CDDP is likely to bind directly to and covalently react with those potent nucleophilic residues of Nrf1 and Nrf2, so that the resulting changes in the biotopological conformations and subcellular tempospatial localizations of these two CNC-bZIP factors affect their protein stability and transcriptional activity. Such stability and activity of both Nrf1 and Nrf2 can also be induced or enhanced in response to oxidative stress triggered by CDDP (Figure 1L). All that is true, a series of the experimental evidence has been provided in this study, demonstrating distinct activation of Nrf1 and Nrf2 by CDDP, as accompanied by discrepant protein stability and transcriptional activity of both factors to regulate the relevant signaling responses.

Interestingly, although CDDP caused significant increases in the expression of Nrf1 and Nrf2 at their mRNA and protein levels in *WT* cells, by contrast, basal and CDDP-stimulated abundances of Nrf1 were also markedly down-regulated by loss of *Nrf2^−/−^*, aside from being almost completely abolished by specific knockout of *Nrf1α^−/−^*. Conversely, the loss of *Nrf1α* resulted in a striking augmentation of basal Nrf2 expression, but its CDDP-inducible expression was remarkably down-regulated in *Nrf1α^−/−^* cells, particularly at its mRNA levels. Such inter-regulation of between Nrf1α and Nrf2 at distinct layers indicate there exists a bi-directional positive and negative feedback circuit tightly monitoring the gene transcription of both CNC-bZIP factors to their protein expression levels. This is due to the fact that the proteasomal dysfunction resulting from *Nrf1α^−/−^* cells leads to a basal accumulation of hyperactive Nrf2 under the normal conditions, which is also placed within a positive feedback loop so to enhance transcriptional expression of its own gene *Nfe2l2 per se*. In other words, such basal protein expression of Nrf2 is regulated negatively by Nrf1α, but CDDP-inducible transcriptional expression of Nrf2 appears to be incremented by Nrf1α, whereas the latter Nrf1α expression is also, in turn, positively regulated by Nrf2 under both basal and CDDP-stimulated conditions. Of note, the remnant Nrf1α factor in *Nrf2^−/−^* cells was still induced by CDDP, whilst a biphasic expression pattern of Nrf2 in CDDP-stimulated *Nrf1α^−/−^*cells was distinctive from its protein expression in *WT* cells. This finding suggests that Nrf1α and Nrf2 are also likely monitored by as-yet-unidentified regulators, besides themselves, particularly during intervention with CDDP. However, it is of crucial importance to note that this drug intervention causes a remarkably prolonged half-life of the processed Nrf1α-C/D isoforms, whereas the stability of intact full-length Nrf1α-A/B and Nrf2 proteins was only modestly enhanced by CDDP.

A significant increase in the intracellular ROS levels was caused by CDDP after intervention of *WT* cells for 12 h, and then reduced to a considerably lower extent, which was rather higher than its baseline level. Such changes in CDDP-stimulated ROS levels were almost completely abolished in *Nrf1α^−/−^* or *Nrf2^−/−^* cell lines, albeit their basal ROS were intrinsically augmented by the loss of *Nrf1α* or *Nrf2*, when compared to those measured from *WT* cells. Intriguingly, largely similar results of total antioxidant capacity (T-AOC) were also obtained from CDDP-stimulated *WT*, *Nrf1α^−/−^* and *Nrf2^−/−^* cell lines, except that the basal T-AOC was down-regulated by specific loss of *Nrf1α*, but not of *Nrf2*. These data imply a controllable redox balance between CDDP-induced ROS and T-AOC levels varying within distinct cellular responses to this drug, and its maintenance is required for the presence of both *Nrf1α* and *Nrf2*, in order to tightly monitor the intracellular ROS production and elimination. That’s just a real fact that has been corroborated by further experiments revealing distinct regulation of those key redox signaling molecules by Nrf1α and Nrf2 alone or both together. Such redox regulation appears to be dictated primarily by CDDP-inducible expression abundances of *HO-1, GCLC, GCLM, GSR, GLS, SLC7A11 (xCT), CAT, PRDX1, PRDX1, SOD1, SOD2, IDH2, IDH2, TIGAR, ME1, TXN1, TXN2, SRXN1, SESN1* and *SESN3*. Most of these genes regulated by Nrf1 and/or Nrf2 are identified to contain more than one consensus AREs/EpREs sequence in their promoter regions [2, 6].

### 4.2 Distinct contributions of Nrf1 and Nrf2 to the anticancer efficacy of CDDP and its resistance

In this study, we have further provided the experimental evidence, unravelling that CDDP intervention results in a rapid death of *WT* cells, along with a considerably lower IC_50_ (15.6 μM) of its viability, when compared with those of *Nrf1α^−/−^*and *Nrf2^−/−^* cell lines (more than 50 μM). This is due to the potent genotoxicity and cytotoxicity of CDDP resulting from its direct reactional and indirect oxidative damages to DNA, RNA and other biological macromolecules in *WT* cells, leading to severe extents of cell death [26, 29, 33], albeit its apoptosis is only marginally increased. By striking contrast, the loss of *Nrf1α^−/−^* or *Nrf2^−/−^*leads to a substantial tolerance or resistance to this drug intervention of their deficient cells. Furtherly, xenograft model animal experiments uncovered that CDDP caused significant repression of *in vivo* malgrowth of *WT-* rather than *Nrf1α^−/−^-*derived tumors in nude mice, but part of the lagged recovery from such tumor-repressing effect of this drug occurred only in the case of *Nrf1α^−/−^*, even though the hyperactive Nrf2 is accumulated in the former *Nrf1α-*deficient cells. This demonstrates distinct contributions of Nrf1α and Nrf2 to the anticancer efficacy of CDDP in hepatoma cells and its drug resistance.

Further examinations also revealed obvious increases in the expression of H2XA (as a key player in the DNA repair) and its phosphorylated γ-H2XA (as a critical entry marker of the DNA damage response) in CDDP-treated *WT* cells. By contrast, basal and CDDP-stimulated expression abundances of H2XA and γ-H2XA were substantially diminished by specific knockout of *Nrf1α^−/−^*, but conversely augmented by the loss of *Nrf2^−/−^*. By scrutinizing the CHIP-sequencing data obtained from (www.encodeproject.org/experiments/ENCSR543SBE or ENCSR488EES), it was, to our surprise, found that a remarkable binding activity of Nrf1, rather than Nrf2, to the promoter regions of *H2XA,* and other key genes *XPA, TOP2A* and *MSH2*, is likely executed in putative DNA damage repair response to CDDP, implying they are the most proper cognate targets of Nrf1, but not Nrf2. Such decreased expression of *H2XA* in *Nrf1α^−/−^* cells was accompanied by down-regulation of *POLH, MSH2, TOP2A, ATM, TP53,* and *BCL2* (as a pro-apoptotic factor), but also with up-regulation of *XPA, REV3L, BRCA3, GSTO3* (required for the metabolism of xenobiotics and carcinogens), *ABCB1* (encoding P-glycoprotein, as a member of the multidrug resistance family), except for Nrf2. However, the opposite expression trends of such genes are determined exactly in *Nrf2^−/−^* cells. Altogether, these indicate distinct roles of Nrf1α and Nrf2 in regulating the DNA damage repair, anti-apoptotic, antioxidant, and even resistance responses to CDDP. Such being the case, it is inferable that that Nrf1α appears to be an indispensable regulator of the DNA repair, whilst Nrf2 serves as a key player in the resistance response. Thereby, the resulting defective DNA repair in *Nrf1α^−/−^* cells should result in the genomic instability insomuch as to promote its malignant transformation, whereas hyperactive Nrf2 enables resistance of *Nrf1α^−/−^*tumors to this drug intervention.

The latter notion is also supported by further experiments of chemoresistance to CDDP, revealing that basal expression abundances of both P-gp and Nrf2 were substantially augmented by a long-term intervention of *WT* hepatoma cells with CCDP for 102 days, while Nrf1 and other examined proteins such as H2XA, γ-H2XA, p53, Bcl2, GPX1 and GSR were only marginally altered or even down-regulated in such intervention conditions. Yet, H2XA degradation induced by chronic low-grade oxidative stress in *Nrf2*-deficient cells were shown to enable the drug chemosensitivity of CDDP to be enhanced in breast cancer patients [56]. In this study of *Nrf2^−/−^*cells, enhanced expression of H2XA and γ-H2XA was accompanied by down-regulation of other key proteins required for the drug resistance and DNA repair response to CDDP. In fact, a number of previous studies have also reported that Nrf2 is indeed required for the resistance to CDDP by up-regulating those drug-metabolizing enzymes and transporters (e.g., P-gp), as well as by enhancing antioxidant, detoxification and cytoprotective defense against CDDP-induced stress [55, 57, 58]. Such hyperactive Nrf2 and enhanced antioxidant capability in *Nrf1α^−/−^*cells enable the deficient tumors to resist against CDDP, even though there exist severe intrinsic oxidative stress and diminished expression of H2XA and γ-H2XA. Altogether, these demonstrate that distinct mechanisms account for the chemoresistance of *Nrf1α^−/−^* rather than *Nrf2^−/−^*-derived cancers to CDDP intervention, albeit the details remain to be elucidated.

### 4.3 Distinct mechanisms by which Nrf1 and Nrf2 contribute to various cellular responses to CDDP

Besides the above-mentioned genotoxic and oxidative stresses, certain proteotoxic stress was also triggered by CDDP in *WT* cells. This is due to the capability of CDDP and its electrophilic derivatives that can directly react with those nucleophilic residues of proteins and other macromolecules, in addition to DNAs/RNAs, and are also internalized in those subcellular organelles (such as endoplasmic reticulum, mitochondria and lysosomes) [55]. Once the homeostasis of such vital organelles and functional proteins was impaired, the consequence leads to particularly proteotoxic stress and unfolded protein responses in the endoplasmic reticulum and mitochondria (i.e., defined as UPR^ER^ and UPR^Mito^, respectively), which may negatively affect the anticancer efficacy of CDDP. In this study, we have provided a series of the experimental evidence, demonstrating distinct involvements of Nrf1α and Nrf2 in mediating diverse signaling responses (including UPR^ER^ and UPR^Mito^) to CDDP.

At the transcriptional mRNA levels measured by transcriptomic sequencing and real-time qPCR, *IRE1, CHOP, FGF21, FOS, PTEN,* and *CDH2* were substantially up-regulated by CDDP in *WT* cells, while *ATF6, Jun, NF-κB,* and *TP53* were only modestly increased, but most of other examined genes *PDI, BIP, PERK, eIF2α, ATF4, XBP1, HSP60, SNAIL1* and *MMP9* were significantly down-regulated by this drug. Such changed expression profiling of these genes responsible for multiple signaling networks is likely to affect the anticancer efficacy of CDDP on hepatoma cells and even its survival adaption to diverse stresses triggered by this drug (Figure 9G). By contrast, basal mRNA expression levels of *PDI, BIP, PERK, eIF2α, ATF4, XBP1, CHOP, HSP60, NF-κB* and *SNAIL1* were substantially down-regulated by the loss of *Nrf1α^−/−^*or *Nrf2^−/−^*. Among them, *CHOP* and *NF-κB* were significantly induced by CDDP in *Nrf2^−/−^* cells, but with only less or no induction of both genes in CDDP-treated *Nrf1α^−/−^* cells. Conversely, *IRE1, FOS, JUN, PI3K, CDH2* and *MMP9* were basally enhanced in *Nrf1α^−/−^* cells, of which *FOS, JUN* and *CDH2* was further augmented by CDDP, while *IRE1, PI3K* and *MMP9* were remarkably reduced by this drug to considerably lower extents than the basal levels measured from *WT* cells. Besides, *PTEN* and *TP53* were basally down-regulated by loss of *Nrf1α^−/−^*, but remained to be strikingly induced by CDDP in this deficient cell line. In *Nrf2^−/−^*cells, although basal *IRE1* and *FGF21* expression was respectively decreased or increased, both genes were substantially induced by CDDP. In addition, no matter what basal expression levels of *FOS, JUN, PI3K, AKT, PTEN, CDH1* and *CDH2* was reduced or enhanced, their inducible expression appeared to be roughly unaffected by this drug. Together, these indicate distinct requirements of Nrf1α and Nrf2 for mediating different signaling responses to CDDP. Of note, the PERK-eIF2α-ATF4 signaling proteins were down-regulated dominantly in *Nrf1α^−/−^* cells, but *Nrf2^−/−^*cells gave rise to increased expression of CHOP as accompanied by increased FGF21 abundance in response to CDDP.

From the above differences, it should also be noted that the protein turnover of those signaling molecules was mostly monitored by Nrf1-targeting proteasomal proteolytic degradation pathways (as described previously [59]), aside from their transcriptional regulation by Nrf1 and Nrf2 alone or together. Thereby, it is inferable that differential or coordinate effects of Nrf1α and Nrf2 are likely exerted on a vast variety of cognate target genes to be regulated at distinct layers from such gene transcription to their protein expression in the response to CDDP. For example, the MAPK signaling pathways were differentially stimulated by CDDP in *WT* cells, but signaling of p-JNK to JUN was enhanced predominantly by this drug in *Nrf1α^−/−^* cells, the MEK-p38 signaling was activated primarily in *Nrf2^−/−^* cells, whilst the ERK signaling was diminished or even abolished by this drug intervention in all three examined cell lines. More interestingly, almost complete abolishment of PTEN in *Nrf1α^−/−^* cells leads to the PI3K-AKT signaling activation, as accompanied by increased abundances of CDH2 and ITGB4 (involved in the EMT process), but just their opposite expression trends were determined in *Nrf2^−/−^* cells. Such distinct deficient effects on these key signaling proteins were also observed in CDDP-stimulated conditions. Collectively, these indicate that the hyperactive Nrf2 in *Nrf1α^−/−^* cells is likely involved in activation of the JNK-JUN and/or PI3K-AKT signaling pathways towards CDH2 or ITGB4, and can also enable the remnant PTEN and p53 responses to CDDP. Moreover, the p38-SNAIL1 signaling induced by CDDP occurred primarily in *Nrf2^−/−^* cells, but the SNAIL1 expression was only marginally enhanced by this drug in *Nrf1α^−/−^*cells, when compared with its equivalent of *WT* cells. This implies a possibility that the MEK-p38 signaling to SNAIL1 activation is much likely mediated by the remnant Nrf1 factor in *Nrf2^−/−^* cells, but not in *Nrf1α^−/−^* cells.

Notably, our previous studies revealed that the hyperactive Nrf2 in *Nrf1α^−/−^* cells promotes glycolysis and lipid biosynthesis, but resulting in more ROS production in the mitochondria, especially under such a higher glycolysis flow pressure [35]. In this study, it was also found that CDDP enabled promotion of glycolysis by up-regulating *GLUT1, HK1, HK2* and *LDHA* in *Nrf1α^−/−^* cells, but the fatty acid metabolism (*CPT1, CPT2, FASN, SCD* and *SREBP2*) was alleviated dominantly in CDDP-intervened *Nrf2^−/−^*cells. Such distinct roles of Nrf1 and Nrf2 in reprogramming of cancer cell metabolism (besides the Warburg’s effect) were further evaluated by untargeted metabolomes. As expected, 896 of differential metabolites with two opposite expression trends in between *Nrf1α^−/−^* and *Nrf2^−/−^* cell lines were selected and then subjected to enrichment of the KEGG pathways. Among them, the first place is taken by the metabolic pathway of *D*-glutamine and *D*-glutamate along with the essential *L*-glutamine. The latter *L*-glutamine was expressed as the first highest metabolite in *Nrf1α^−/−^* cells and converted into the second highest metabolite of 2-oxo-glutarate (as a key complement to the TCA cycle linked to the electron transport chain with the ROS production in the mitochondria). Further examination revealed that the loss of *Nrf1α* caused significant augmentations in basal and CDDP-inducible expression of *GLUL,* a key gene required for the metabolic conversion of *L*-glutamate into *L*-glutamine to yield massive 2-oxo-glutarate. This is held as a main resource to continuously replenish the TCA cycle and accelerate gluconeogenesis and glutathione biosynthesis, in order to provide certain energy and nutrients so to meet the needs of proliferating cancer at a redox steady-state. Such redox metabolism related to NADP^+^/NADPH is likely reprogrammed by CDDP intervention and varied with the loss of *Nrf1α* or *Nrf2*.

The reprogramming of cancer metabolism was further analyzed by metabolome of xenograft tumor tissues, unraveling a main predominant place is taken through the tryptophan metabolism monitored by Nrf1α. In this metabolism pathway, the Kyn branch is required for a *de novo* NAD^+^ biosynthesis. This is because both KYNU and HAAO enzymes catalyze the production of 2-amino-3-carboxylmuconate semialdehyde to enter the nicotinic acid metabolism pathway and hence generate NAD^+^ and other intermediates under the action of QPRT, consequently initiating the NAD^+^ remediation plan (Figure 7C). The relevant catalytic enzymes IDO2, AFMID, and KMO were up-regulated in *Nrf1α^−/−^* tumor tissues, resulting in a higher demand for metabolites 3-Hydroxy-L-kynurenine and 2-amino-3-carboxylic acid semialdehyde, as well as the increased production of NAD^+^ and NADP^+^ to be 1.7 times higher than those measured in *WT* cells. However, an inhibitory effect of CDDP on those catalytic enzymes (e.g., IDO2, KYN, HAAO) was exerted in *Nrf1α^−/−^* tumors, but it remained to be higher than their counterparts of *WT*, implying that these changes are attributable to the expression alteration of Nrf2 to a roughly similar trend. Such nuanced mechanisms could provide an explanation of the better anticancer effect of CDDP on *WT*-tumors than on *Nrf1α^−/−^* tumors (due to hyperactive Nrf2 retained for its resistance to this drug). Overall, it inferable that the anticancer pharmacological effect of CDDP is more likely to depend on the presence of Nrf1 rather than Nrf2, principally by targeting its specific transcriptome, proteome and metabolome.

## 5. Concluding remarks

Since cell fate decision and transformation (during carcinogenesis and malignant progression) are dominantly dictated by distinct sets of those differential expression genes regulated by a limited number of key transcription factors, they hence facilitate to be paved as new anticancer drug targets, as proposed by Darnell *et al* [60] and Lee *et al* [61]. So far, it has been proved that about 30 of transcription factors possess potent tumor-repressing or tumor-promoting effects on the development of cancers by governing transcription of their specific target genes. Of note, dual opposing roles of Nrf2 in promoting or repressing cancers are also accompanied by drug resistance to chemotherapy [58]. By contrast, Nrf1 is endowed with an intrinsic tumor-repressing activity to be executed by its target proteasomal inhibition by bortezomib (a specific inhibitor of core proteasomal enzymatic sites), leading to the selective processing of this membrane-bound protein to yield mature CNC-bZIP factor [62], which provides a theoretical basis for target treatment of breast cancer. Here, we have discovered distinct contributions of Nrf1 and Nrf2 to the anticancer efficacy of CDDP on hepatoma cells and xenograft tumors, and even drug resistance. Such anticancer effect of CDDP is exerted more dependently on the presence of Nrf1 in *WT* tumors, rather than its absence in *Nrf1α^−/−^* tumors (due to the hyperactive Nrf2 retained responsible for its resistance to this drug). More interestingly, the remnant Nrf1 expression in *Nrf2^−/−^*cells can still enable up-regulation of those key genes involved in the DNA damage repair, particularly during CDDP intervention. Furtherly, a series of experiments and big-data analyses demonstrated distinct cellular metabolisms, molecular pathways and signaling mechanisms by which Nrf1 and Nrf2 differentially contribute to the tumor-repressing effect of CDDP by targeting their specific transcriptomes, proteomes and metabolomes. The putative targets of Nrf1 and/or Nrf2 are likely responsible for multi-hierarchal signaling pathways to cognate gene regulatory networks, in order to enable the reprogramming of cancer cell metabolism and even cell reshaping in the adaptive responses to CDDP (Figure 9G, and also see the Graphical Abstract). Overall, this study has laid a solid foundation for further exploration of Nrf1 as a drug target, and also provides a new strategic direction for the future translational research on its targeted activators

## Supporting information

Supplement materials related results of experiment

## Author contributions

R.W. designed and performed most of the experiments, collected all the relevant data, and wrote a draft of this manuscript with most figures and supplemental information. J.F and K.L. performed the subcutaneous xenograft tumor experiments and some of drug intervention. Both S.H. and M.W. helped R.W. with mice anatomy, tissue section, metabolomics and transcriptome data analysis and other experiments. Both F.C. and S.L. helped R.W performed some of the western blotting and data collection. Lastly, Y.Z. designed and supervised this study, analyzed all the data, helped to prepare all figures with cartoons, rewrote and revised the paper. All these co-authors have read and agreed to the published version of the manuscript.

## Supplementary Materials

The supporting information, including three supplemental figures and also one supplemental tables.

## Ethics statement

The animal study was reviewed and approved by the University Laboratory Animal Welfare and Ethics Committee (with institutional licenses SCXK-PLA-20210211).

## Acknowledgments

We are greatly thankful to Dr. Yonggang Ren (North Sichuan Medical College, Sichuan, China) and Dr. Lu Qiu (Zhengzhou University, Henan, China) for their involvement in establishing the indicated cell lines used in this study, and also thank to all other present and past members of Zhang’s laboratory for giving critical discussion and invaluable help with this work. This study was funded by the National Natural Science Foundation of China (NSFC with two projects 82073079 and 81872336) awarded to Prof. Yiguo Zhang (at Chongqing University). This is also, at part, supported by the Initiative Foundation of Jiangjin Hospital affiliated to Chongqing University (2022qdjfxm001).

