## Supplement materials related results of experiment for "Distinct mechanisms by which antioxidant transcription factors Nrf1 and Nrf2 as drug targets contribute to the anticancer efficacy of Cisplatin on hepatoma cells"

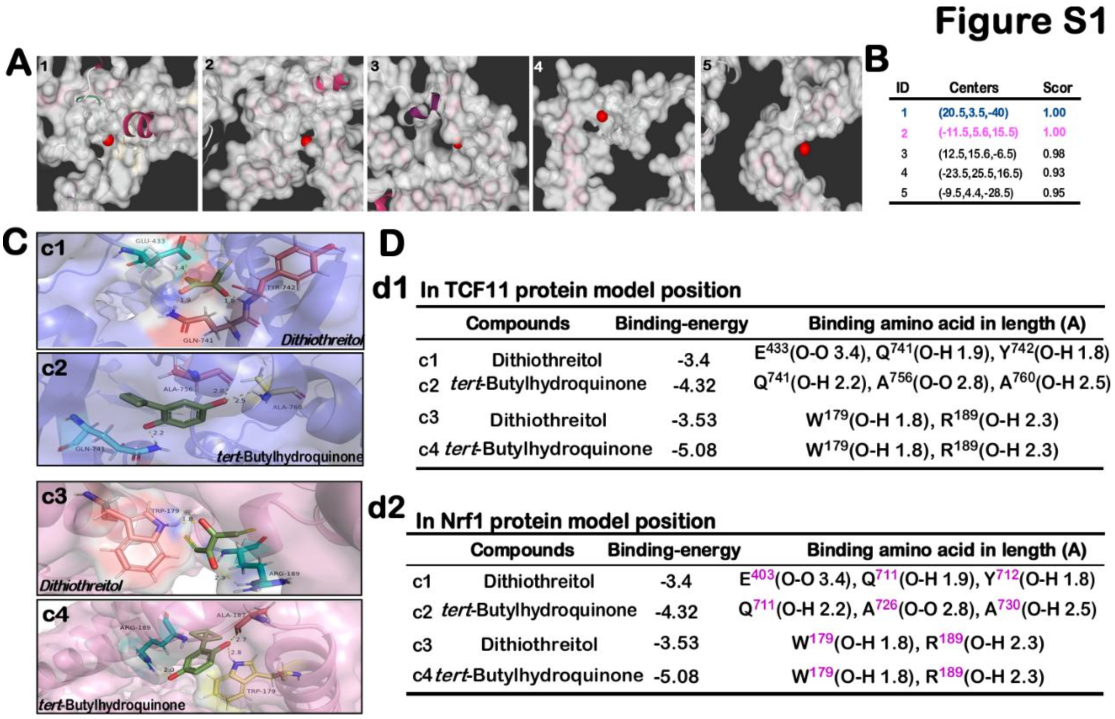


**Figure S1. Prediction of the active site of *Nrf1/TCF11* protein and docking with *tBHQ* and DTT molecules. (A)** The Deepsite online tool (https://www.playmolecule.com/deepsite/ ) was used to predict the active site in the Nrf1/TCF11 protein, and the top five site coordinates (*x, y, z*) **(B)** were obtained. The two coordinate values with the highest scores were selected for virtual molecular docking with antioxidant *tBHQ* **(c1, c3)** and reducing stressor DTT **(c2, c4)** through AUTODOCK 4.0. Based on the obtained docking information, label such as docking site amino acids and bond lengths in PyMOL. For important details of binding energy, docking amino acid, bond length and so on, noted as the TCF11 **(d1)** and Nrf1 **(d2)** respectively through tables **(D)**.

### Figure S2

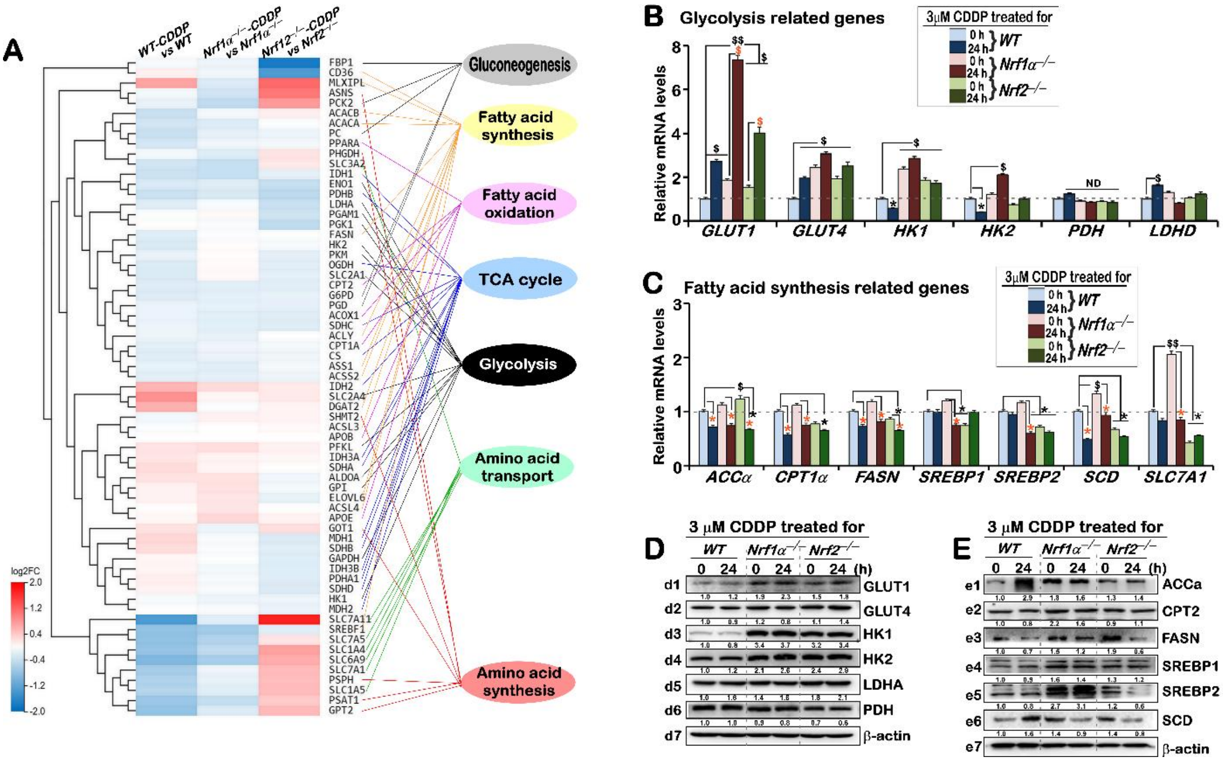


**Figure S2. Distinction expression of glycolysis and fatty acid biosynthesis related genes in the different genotypes cells after CDDP intervention. (A)** Differential expression heat-map of glycose, fatty acid metabolism and amino acid transported related genes in transcriptome in *WT, Nrf1α*$^{-/-}$ and *Nrf2*$^{-/-}$ cell lines after intervention with 3 μM CDDP. **(B,C)** The regulatory effects of 3 μM CDDP on those glycolysis of genes *GLUT1, GLUT4, HK1, HK2, PDH, LDHD* and fatty acid metabolism related genes (*ACCα, CPT1, FASN, SREBP1, SREBP2, SCD, SLC7A1*) in distinct genotypic cell lines after 0 and 24 h of intervention, respectively. The real-time qPCR data were shown as fold changes (mean ± SD, n = 3 × 3), which are representative of at least three independent experiments being each performed in triplicates. Significant increases ($, p < 0.05; $$, p < 0.01) and significant decreases (*$p < 0.05$) were statistically analyzed when compared with WT (measured at 0 h). In addition, some symbols "$ or *" indicate significant differences in *Nrf1α*$^{-/-}$ or *Nrf2*$^{-/-}$ cells compared to their respective t0 after CDDP intervention. **(D,E)** Most of their related proteins of GLUT1 **(d1)**, GLUT4 **(d2)**, HK1 **(d3)**, HK2 **(d4)**, PDH **(d5)**, LDHD **(d6)**, ACCα **(e1)**, CPT1 **(e2)**, FASN **(e3)**, SREBP1 **(e4)**, SREBP2 **(e5)**, and SCD **(e6)** were determined by Western blotting with distinct antibodies . The intensity of those immunoblots was then quantified by the Quantity One 4.5.2 software, and shown below the relevant protein bands.

**Figure S3**

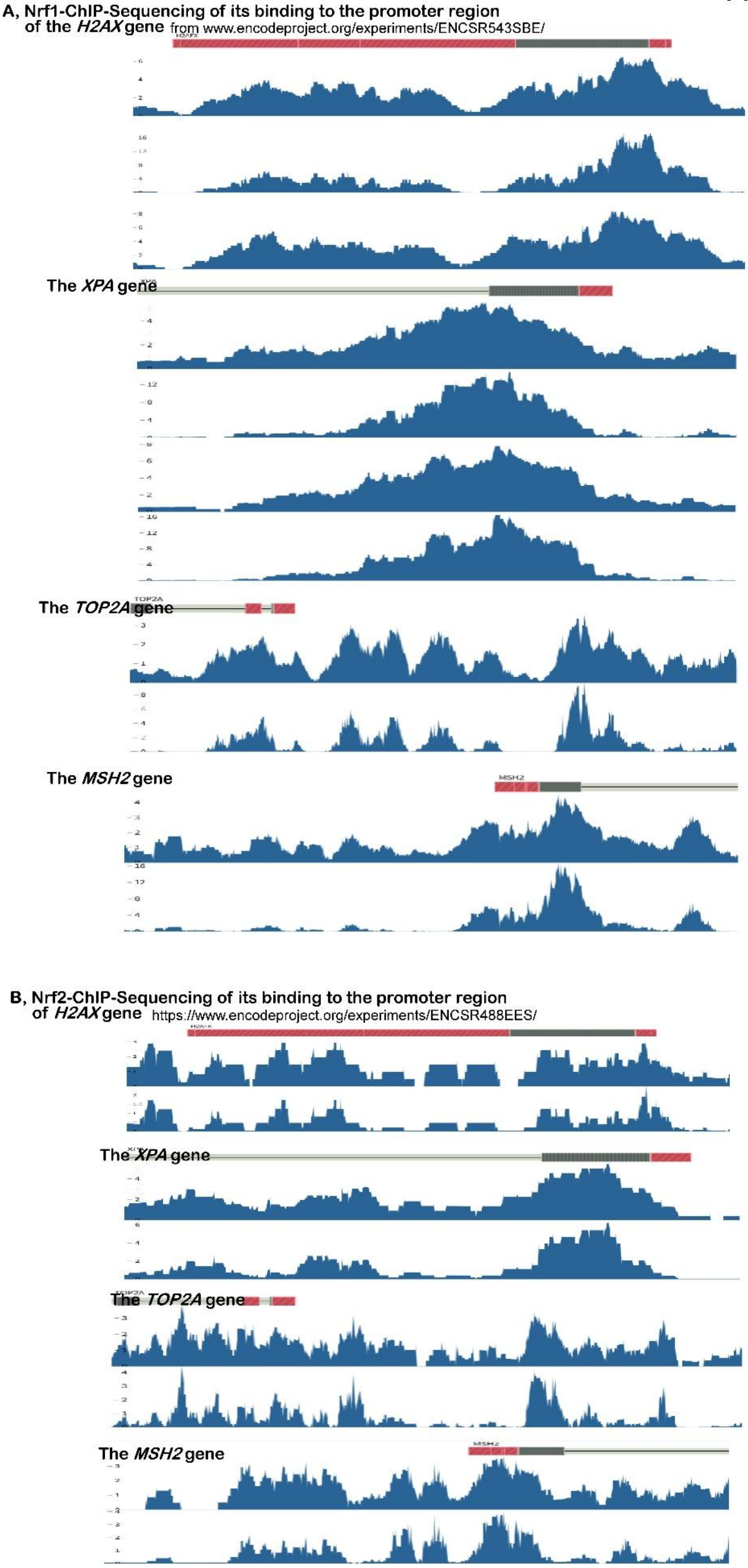


**Figure S3. Chromatin immunoprecipitation (ChIP)-sequencing mediated by Nrf1 and Nrf2, the data of which are obtained from** www.encodeproject.org/experiments/ENCSR543SBE or ENCSR488EES
